# The DAXX-SREBP axis promotes oncogenic lipogenesis and tumorigenesis

**DOI:** 10.1101/2020.12.31.424997

**Authors:** Iqbal Mahmud, Guimei Tian, Jia Wang, Jessica Lewis, Aaron Waddell, McKenzie L. Lydon, Lisa Y. Zhao, Jian-Liang Li, Hamsa Thayele Purayil, Zhiguang Huo, Yehia Daaka, Timothy J. Garrett, Daiqing Liao

## Abstract

De novo lipogenesis produces lipids for membrane biosynthesis and cell signaling. Elevated lipogenesis is a major metabolic feature in cancer cells. In breast and other cancer types, genes involved in lipogenesis are highly upregulated, but the mechanisms that control their expression remain poorly understood. DAXX modulates gene expression through binding to diverse transcription factors although the functional impact of these diverse interactions remains to be defined. Our recent analysis indicates that DAXX is overexpressed in diverse cancer types. However, mechanisms underlying DAXX’s oncogenic function remains elusive. Using global integrated transcriptomic and lipidomic analyses, we show that DAXX plays a key role in lipid metabolism. DAXX depletion attenuates, while its overexpression enhances, lipogenic gene expression, lipid synthesis and tumor growth. Mechanistically, DAXX interacts with SREBP1 and SREBP2 and activates SREBP-mediated transcription. DAXX associates with lipogenic gene promoters through SREBPs. Underscoring the critical roles for the DAXX-SREBP interaction for lipogenesis, SREBP2 knockdown attenuates tumor growth in cells with DAXX overexpression, and a DAXX mutant unable to bind SREBPs are incapable of promoting lipogenesis and tumor growth. Our results identify the DAXX-SREBP axis as an important pathway for tumorigenesis.

## INTRODUCTION

Cancer cells exhibit elevated de novo intracellular lipogenesis, resulting in increased levels of fatty acids, membrane phospholipids, and cholesterol (1). Notably, de novo lipogenesis contributes minimally to the overall lipid content of normal non-proliferating cells, which generally rely on the uptake of lipids from the circulation. In contrast, highly proliferative cancer cells show strong avidity to acquire elevated lipids and cholesterol through either enhancing the uptake of exogenous (or dietary) lipids and lipoproteins or hyperactivating their endogenous de novo lipid synthesis mechanism (1,2). Increased de novo lipogenesis in cancer cells is thought to supply lipids for the synthesis of membranes and signaling molecules during rapid cell proliferation and tumor growth, due to limited availability of lipids from the circulation in the tumor microenvironment (1,3). De novo lipogenesis is controlled by several transcription factors, such as the sterol regulatory element-binding proteins, SREBP1 and SREBP2 (SREBP1/2), that have been shown to play an important role in maintaining lipid synthesis in cancer (4). SREBP1/2 precursors are sequestered in endoplasmic reticulum. When sterol supply is low, SREBP1/2 are transported to the Golgi apparatus where they are cleaved by proteases, and the N-terminal domains of SREBPs are then released and imported into the nucleus to promote transcription of genes that contain the sterol regulatory elements (*SREs*) required for lipogenesis.

Independently of intracellular lipid levels, oncogenic drivers, including KRAS and PI3K, promote de novo lipogenesis in BC and other cancer types converging on mTORC1 activation (1,5-7). mTORC1 promotes S6K1-dependent SREBP1/2 processing (8). The phosphatidate phosphatase Lipin-1 sequesters mature SREBP1/2 in the nuclear lamina, thereby preventing SREBP1/2 from activating gene expression. mTORC1 directly phosphorylates Lipin-1, which inhibits its nuclear translocation and thus restores SREBP activity (9). mTOR signaling also indirectly stabilizes SREBP1/2 by opposing phosphorylation-dependent ubiquitination of SREBP1/2 by the E3 ubiquitin ligase FBXW7 and subsequent proteasomal degradation (10-12). Notably, tumors efficiently convert acetate to acetyl-CoA (13), which is predominantly used for lipid synthesis (14), highlighting the need for cancer cells to activate lipogenic enzymes (15). While the dependence on de novo lipogenesis in cancer is well documented, the mechanisms that control SREBP-mediated transcription underlying oncogenic de novo lipogenesis remain poorly understood.

DAXX, originally discovered as a context-dependent regulator of cell death or survival (16-18), has an extensively documented role in transcription regulation through interacting with transcription factors including p53 (19) and NF-κB (20). More recent studies have defined DAXX as a specific chaperone for the histone variant H3.3 (21-23). DAXX binds specifically to the H3.3/H4 dimer and deposits it onto chromatin (24,25). Emerging evidence suggests that DAXX has an oncogenic role in diverse cancer types (26,27), which appears to be linked to its functions in gene regulation (18,27,28). Whereas the levels of DAXX expression directly correlate with its ability to promote tumor growth (18,26-28), the molecular mechanisms underlying DAXX’s oncogenic function remain to be defined.

In this study, we identified DAXX as a novel regulator of oncogenic lipogenesis through its interaction with SREBP1/2, leading to activating lipogenic gene expression programs and the promotion of cancer cell proliferation in vitro and tumor growth in vivo. Our studies define the DAXX-SREBP axis as a previously unrecognized oncogenic pathway.

## MATERIALS AND METHODS

### Cell culture

Cell lines used for this study were obtained from ATCC (Manassas, VA) and authenticated by Genetica DNA Laboratories (Burlington, NC). Cells were cultured in Dulbecco’s Modified Eagle’s Medium (DMEM with 4.5 g/L glucose, L-glutamine and sodium pyruvate, Corning, Tewksbury, MA) with 10% bovine calf serum (HyClone, GE Healthcare Bio-Sciences, Pittsburgh, PA), penicillin (10 units/mL), and streptomycin (10 µg/mL) (the complete DMEM medium). The T47D cell line was cultured in DMEM plus 10% fetal bovine serum (Atlanta Biologics, Atlanta, GA), penicillin (10 units/mL), and streptomycin (10 µg/mL). To culture cells in serum starvation condition, serum-containing medium was removed from cell cultures after overnight culture and the culture was washed once with phosphate-buffered saline (PBS, without calcium and magnesium, Corning). Cells were then cultured in serum-free DMEM. For culturing cells in suspension (3D culture), plates were coated with a 1:1 mixture of Matrigel (Corning) and complete DMEM medium. A desirable number of cells were suspended in the Matrigel and medium mixture and layered on the top of the solidified Matrigel. Complete DMEM medium was added after the Matrigel was solidified. Medium was replaced with fresh complete medium every three days. Colonies were imaged under a microscope; colony numbers and sizes were quantified.

### DNA constructs

cDNAs for wild-type (WT) DAXX and mutants with a 5’ coding sequence for the FLAG epitope tag and a 3’ coding sequences for the MYC and 6x His tags were cloned into a lentiviral vector under the control of the cytomegalovirus immediate early (CMV *IE*) promoter. GFP-DAXX constructs were cloned in the pEGFP-C2 vector. A short hairpin RNA (shRNA) targeting the DAXX coding sequence (nucleotide 624-642, 5’-GGAGTTGGATCTCTCAGAA-3’) was cloned into a lentiviral vector under the control of the human H1 promoter. An shRNA construct with a scrambled sequence (Plasmid # 36311) was from Addgene. Expression vectors for mature SREBP1a (Plasmid # 26801), mature SREBP1c (Plasmid # 26802), and mature SREBP2 (Plasmid # 26807) were purchased from Addgene. The shRNA clones for *SREBF1* (TRCN0000020607 and TRCN0000020605), and *SREBF2* (TRCN0000020667 and TRCN0000020668) were from the human pLKO.1 TRC Library collection at the University of Florida. The *SREBF2* shRNA vector TRCN0000020667 was used to knockdown *SREBF2* expression in MDA-MB-231 cells with DAXX OE. A *SREBF2* promoter fragment was PCR amplified from the genomic DNA isolated from MDA-MB-231 cell line and cloned at sites upstream of the firefly luciferase reporter by the Gibson assembly method. The DNA sequence was confirmed by Sanger sequencing. The PCR primers are shown in Supplementary Table S1. Stable expression of cDNA and shRNA was established through lentiviral transduction of cell lines and puromycin (2 µg/mL) selection. The derived cell lines were cultured with DMEM without puromycin.

### Microarray, RNA-seq and qRT-PCR

Cells were cultured in the complete DMEM or serum-free DMEM, and total RNAs were isolated using the RNeasy kit (Qiagen) for microarray and RNA-seq analysis. For microarray experiments, the RNAs were then processed for microarray hybridization to the Affymetrix GeneChip Human Transcriptome Array 2.0 as described previously (29,30). Total RNAs were used for RNA-seq library constructions and sequencing was done with 20M raw reads/sample using the Illumina Platform PE150 at Novogene Corporation Inc. (Sacramento, CA).

For quantitative real-time PCR (RT-qPCR), the isolated RNAs were reverse transcribed with random hexamers using 2 µg of total RNA, an RNase inhibitor, and reagents in the Multiscribe reverse transcriptase kit (Life Technologies). The resulting cDNAs were diluted and used as input for qPCR using the SYBR green detection method. The relative levels of gene expression were determined using the ΔΔCt method with the Ct values of ACTB expression as the common normalizer. The primers for qPCR and other applications are shown in Supplementary Table S1.

### Immunoprecipitation (IP) and Immunoblotting

Cell pellets were resuspended in the IP lysis buffer (50 mM Tris-HCl, pH 7.5, 0.5% Igepal-CA630, 5% glycerol, 150 mM NaCl, 1.5 mM MgCl_2_, and 25 mM NaF) containing 100-fold diluted protease inhibitor cocktail (Millipore-Sigma P8340). The cell suspension was subjected to two freezing/thawing cycles. The cell lysates were then centrifuged at 15,000 rpm at 4°C for 20 min. The supernatant was used for IP with a control or an antibody to a specific protein at 2 µg per IP in the presence of protein A-agarose beads. The beads were resuspended in the IP lysis buffer along with one fifth of the volume of the 6x SDS sample buffer (0.375 M Tris-HCl, pH 6.8, 12% SDS, 60% glycerol, 0.6 M DTT, and 0.06% bromophenol blue). Samples were heated at 95°C for 5 min and chilled on ice for 2 min. After brief centrifugation, the samples were loaded on a 4-20% gradient gel (Novex Tris-Glycine Mini Gels, ThermoFisher). Proteins were then electrotransferred to an Immobilon®-P polyvinylidene fluoride (PVDF) membrane (Millipore). Membrane was blocked with 5% non-fat milk, incubated with a primary antibody and a proper secondary antibody. The proteins were detected using a chemiluminescent detection kit (Millipore) and the Fuji Super RX-N X-ray films or an Amersham Imager 680.

For immunoblotting analyses of cell lysates of monolayer cultures, medium was removed from culture plates and 1x Passive Lysis buffer (Promega) was added. The plates were frozen at −80°C overnight and then thawed at room temperature. The lysates were transferred to a centrifuge tube. To prepare tumor lysates, xenograft tumor tissues were fragmented in the presence of liquid nitrogen, approximately 50 mg of tumor fragment was homogenized in 1 mL of 1X RIPA lysis buffer on ice using a micro-homogenizer. After brief sonication at a low power output for 5 sec on ice, the lysates were cleared by centrifugation at 13,000 rpm for 15 min at 4 °C. Protein contents were quantified using a Qubit protein assay kit. Protein extracts from cell culture or tumor lysates were subjected to SDS-PAGE and electro-blotting as above. The antibodies used for this study are listed in Supplementary Table S2.

### Proximity Ligation Assay (PLA)

The PLA reagents were obtained from Millipore-Sigma (DUO92101-1KT). The assays were performed following the manufacturer’s protocol. The antibodies against SREBP2 (Abcam, ab30682), SREBP1 (ProteinTech, 4088-1-AP), DAXX (5G11 hybridoma supernatant) were used for the PLA experiments. The number of PLA signal dots was quantified as described previously (31).

### De novo lipogenesis assays

Cells (0.5 million per well) were plated in a 6-well plate in complete DMEM medium in triplicate. At 24h after seeding, cells were washed once with PBS and cultured in serum-free DMEM for 16 h; 5 µCi of [1-^14^C] acetate (NEC084H001MC, Perkin Elmer, Waltham, MA, USA) per mL was added and the cells were cultured for four more hours. Cells were then washed twice with PBS and trypsinized. Cells were pelleted and resuspended in 0.5 mL of 0.5% Triton X-100. The protein concentration of the lysates was determined for normalization. The lysates were extracted with ice cold chloroform/methanol (2:1 v/v). After centrifugation at 1,000 rpm for 20 min, the organic phase was collected and air dried. The radioactivity was determined with a liquid scintillation counter (Beckman LS 5000TD). The radioactivity was normalized against protein concentration.

### Liquid chromatography (LC)-mass spectrometry (MS) experiments

For lipid analysis, we used these internal lipid standards: triglyceride (TG 15:0/15:0/15:0 and TG 17:0/17:0/17:0, Sigma-Aldrich), lysophosphatidylcholines (LPC, 17:0 and 19:0), phosphatidylcholines (PC, 17:0/17:0 and 19:0/19:0), phosphatidylethanolamines (PE, 15:0/15:0 and 17:0/17:0), phosphatidylserines (PS, 14:0/14:0 and 17:0/17:0), and phosphatidylglycerols (PG, 14:0/14:0 and 17:0/17:0) (Avanti Polar Lipids, Alabaster, AL). The lipid standards were dissolved in 2:1 (v/v) chloroform/methanol to make a 1000 ppm stock solution and a working 100 ppm standard mix was then prepared by diluting the stock solution with the same solvent mixture. For sample normalization, total protein concentration in each sample was determined using a Qubit 3.0 Fluorometer.

Cell lines with a control vector, an shRNA against an indicated gene, WT DAXX, or the del 327-335 mutant were cultured with the complete DMEM. When cells grew to approximately 80% confluency, they were washed twice with PBS and cells were detached using a cell lifter. Cell pellets were washed twice with 40 mM ammonium formate (AF). The cell pellets were resuspended in 50 µL of AF with vortex in a glass vial and subjected to high efficient bead beater cell disruption to release intracellular lipids. A small amount of the homogenized cell pellet was taken for Qubit protein concentration determination. Lipids were extracted by adding ice-cold chloroform (2 mL) and methanol (1 mL) along with 20 µL of internal standard mixtures. The extraction mixture was incubated on ice for 1 h with occasional vortex mixing. Finally, 1 mL H_2_O was added to the mixture, which was incubated for 10 min with occasional vortex mixing. Samples were then centrifuged at 2,000 rpm for 5 min. The lower phase (organic layer) was collected in a separate glass vial and subjected to dry under nitrogen gas at 30 °C using a dryer (MultiVap, Organomation Associates). Dried samples were reconstituted by adding 50 µL isopropyl alcohol and transferred to a glass LC vial with insert. Samples were loaded to an auto-sampler at 5 °C.

For analyzing lipids, we ran samples for quality control (QC) in each instrument run. A pooled QC sample (a 25 µL aliquot) for each extraction was injected after analyzing every five samples. The pooled QC sample was run to assess system reproducibility, and a blank (solvent mixture only) was used to flush the column. We did not observe any changes regarding the number of background ions, which always corresponded to the specific solvent used for lipid extraction. Also, we did not notice any effects on reproducibility of ion source regardless of solvents used for extraction. The stability and repeatability of the instruments were evaluated using identical neat QC samples (a mixture of all internal standards in deuterated form) throughout the process of sample injection. Principal component analysis (PCA) was performed to evaluate the variation of QC samples. All neat QC samples clustered together, confirming the stability and reproducibility of our experimental lipid analysis system.

For data collection, processing, and analysis, we used a Dionex Ultimate 3000 UHPLC system coupled to a Q Exactive™ hybrid quadrupole-orbitrap mass spectrometer operated in HESI-positive and negative ion mode. A Supelco Analytical Titan reverse-phase column (RPC) C18 (2.1 × 75 mm with 1.9 μm monodisperse silica) equilibrated at 30 °C with solvents A (acetonitrile and water 60:40, v/v) and B (isopropyl alcohol, acetonitrile, and water 90:8:2, v/v/v) as mobile phases was used for data collection. The flow rate was 0.5 mL/min, and the injection volume was 5 µL. The total run time was 22 min, including a 2-min equilibration. The MS conditions for positive and negative ion modes were spray voltage at 3.5 kV, sheath gas at 30 arbitrary units, sweep gas at 1 arbitrary unit, auxiliary nitrogen pressure at 5 arbitrary units, capillary temperature at 300 °C, HESI auxiliary gas heater temperature at 350 °C, and S-lens RF at 35 arbitrary units. The instrument was set to acquire in the mass range of most expected cellular lipids and therefore *m/z* 100–1500 was chosen with a mass resolution of 70,000 (defined at *m/z* 200). Global lipid profiling was performed using full scan and ddMS2 (data dependent MS-MS).

Data were recorded from 0.0 to 17 min as total ion chromatography (TIC) and then corresponding MS data were extracted using Thermo Xcalibur (version 2.2.44). After data collection, raw data files were converted to mzXML format using the Proteowizard MSConvert software. MZmine 2.15 (freeware) was used for mass detection with mass detector centroid noise set at 1.0E05 using only MS level 1 data; chromatogram building and deconvolution were then applied (m/z tolerance, 0.005 or 10 ppm; retention time tolerance, 0.2 min; minimum time span, 0.1 min; and minimum height, 5.0E05) followed by isotope grouping, alignment (m/z tolerance, 0.005 or 10 ppm; retention time tolerance, 0.2 min), and gap filling (m/z tolerance, 0.005 or 10 ppm; retention time tolerance, 0.2 min, and intensity tolerance 25%). MZmine-based online metabolite search engine KEGG, MMCD database, XCMS online database, Metaboanalyst 3.0, R program, and internal retention time library were used for the identification and analysis of metabolites.

### In vivo tumor growth

All mice were maintained under pathogen-free conditions. Female NSG (NOD.Cg-Prkdc^scid^Il2rg^tm1Wjl^/SzJ) mice, between the ages of 4-6 weeks, were injected subcutaneously in a mammary fat-pad area with one million cells in 100 µL of complete DMEM (MDA-MB-231-derived cell lines) or in a suspension of 50 µL of Matrigel and 50 µL of cell suspension (MDA-MB-468-derived cell lines). Tumor growth was monitored by measuring tumor dimensions using a digital caliper once a week until endpoint. Tumor volume was calculated with the formula ½ x length x width^2^. At the endpoint, mice were euthanized, tumors were excised, weighted, and photographed. Tumor lysates were prepared for immunoblotting analysis. Animal use has been approved for this project by the University of Florida IACUC.

### Chromatin immunoprecipitation (ChIP)

The panel of MDA-MB-231-derived cell lines (control and WT DAXX OE) were cultured in complete DMEM. ChIP experiments were performed essentially as described (32). Briefly, at about 90% confluency, the cells were crosslinked by adding 37 % formaldehyde to the final concentration of 1% for 10 min at room temperature. Crosslinking was stopped by adding glycine to the final concentration of 125 mM. Cells were lifted, washed with cold PBS, and pelleted by centrifugation. The cells were resuspended in a swelling buffer in the presence of the protease inhibitor cocktail (Sigma) and then pelleted and resuspended in the SDS lysis buffer. The lysates were transferred to a Covaris microTUBE and sonicated with an E220 Covaris Ultrasonicator. Chromatin fragmentation (∼500 bps) was verified by agarose gel electrophoresis. The fragmented chromatins were diluted and incubated with a control IgG and the DAXX mAb (5G11) along with protein A/G magnetic beads. The beads were washed sequentially with a low salt buffer, high salt buffer, LiCl buffer, and TE buffer (twice). The immunoprecipitated chromatins were eluted at 65 °C for 15 min, and the eluted chromatins were subjected to proteinase K digestion at 65 °C for 3 h. The DNAs were recovered through a Qiagen mini-prep column. The immunoprecipitated DNAs were used for qPCR and library construction and high throughput sequencing using an Illumina Hi-Seq 2500 sequencer.

### Bioinformatics analysis

We analyzed gene expression based on publicly available datasets. Gene expression data for normal, benign, primary, and metastatic tumor samples were included for our analysis. Normalized expression levels for specific genes were compared between different sample types. Computations were conducted in R statistical package (https://www.r-project.org/) and in GraphPad Prism 7.0. For Ingenuity Pathway Analysis (IPA), genes that were differentially expressed (fold-change over ±1.3 and p-value < 0.05) were used for the Ingenuity Pathway Analysis (Ingenuity Systems, Qiagen Bioinformatics, http://www.ingenuity.com). Gene Set Enrichment Analysis (GSEA) was performed using the Java desktop software (http://software.broadinstitute.org/gsea/index.jsp), as described previously (33). The GSEA tool was used in pre-ranked mode with all default parameters. For microarray data analysis, probe set files (.cel file) were normalized by RMA algorithm and analyzed using both R statistical package as well as Affymetrix expression and transcriptome console software from ThermoFisher Scientific. For RNA-seq data analysis, we used the RNAseq data analysis pipeline reported previously (34). Briefly, fastq files were aligned to Genome Reference Consortium Human Build 38 (GRCh38) using HISAT2 (35); the transcripts assembling was performed using StringTie (36) with RefSeq as transcripts ID; and the normalized counts (by FPKM) was called using Ballgown (37). The differential expression analysis was performed using R package limma (38); and the pathway enrichment analysis was performed using ingenuity pathway analysis. ChIP-seq sequencing reads (Fastq files) were mapped to the human genome (GRCh37/hg19) using Bowtie2 (39), where option –local was specified to trim or clip unaligned reads from one or both ends of the alignment. Genome browser BedGraph tracks and read density histograms were generated using SeqMINER. Peak finding and annotation to the nearest Refseq gene promoter was performed and de novo motif discovery was carried out using HOMER (40).

### Statistical analysis

Gene expression assays were conducted in two to three biological replicates. Metabolic profiling assays were performed in four to six replicates. Data are presented as the mean along with standard error of the mean (SEM). Student’s t-test was used to compare two groups of independent samples. For all data analysis, p<0.05 was considered statistically significant.

## RESULTS

### Transcriptomic profiling implicates DAXX in promoting lipogenic gene expression

Bioinformatic analyses of clinical BC samples of The Cancer Genome Atlas (TCGA) and the Clinical Proteomic Tumor Analysis Consortium (CPTAC) datasets revealed that DAXX mRNA and protein levels are elevated in all four major BC subtypes with highest levels in the triple-negative BC (TNBC) subtype (Figure 1A and B), suggesting a potential oncogenic role for DAXX (18). To understand the function of DAXX in cancer, we used gain and loss of function approaches: genetically depleted endogenous DAXX or overexpressed wild-type (WT) DAXX in the TNBC cell line MDA-MB-231 (Figure 1C). Transcriptomic analyses using microarray and RNA-seq revealed distinct gene expression profiles for cells with DAXX mRNA knockdown (KD) and WT DAXX overexpression (OE) in comparison to control (CTL) cells (Figure 1D and E). Unexpectedly, Ingenuity Pathway Analysis (IPA) of differentially expressed genes in cells with DAXX KD or OE in comparison to CTL cells revealed a marked downregulation and upregulation, respectively, of the de novo lipogenesis pathways. Lipogenesis regulators (*SREBF1/2* encoding SREBP1/2 and SCAP) were among the most highly inhibited upstream regulators in the KD cells, while WT DAXX OE activated *SREBF1/2* (Figure 1F). Correspondingly, the cholesterol biosynthesis via the mevalonate pathway were among the top canonical pathways identified by IPA (Supplementary Figure S1A). Most of the genes in the biosynthesis of cholesterol, fatty acids, glycerolipid, and glycerophospholipids are affected by DAXX expression levels (Figure 1E). Gene Set Enrichment Analysis (GSEA) of transcriptomic data confirmed suppression and activation of the de novo lipogenesis pathway by DAXX KD and WT OE, respectively (Supplementary Figure S1B and C). Of note, several transcriptional regulators that are known to interact with DAXX such as JUN and PML (18) were also affected by DAXX expression levels. Interestingly, the insulin receptor (INSR) pathway that regulates intracellular lipid production (41) also seems to be positively regulated by DAXX (Figure 1F).

**Figure 1.**
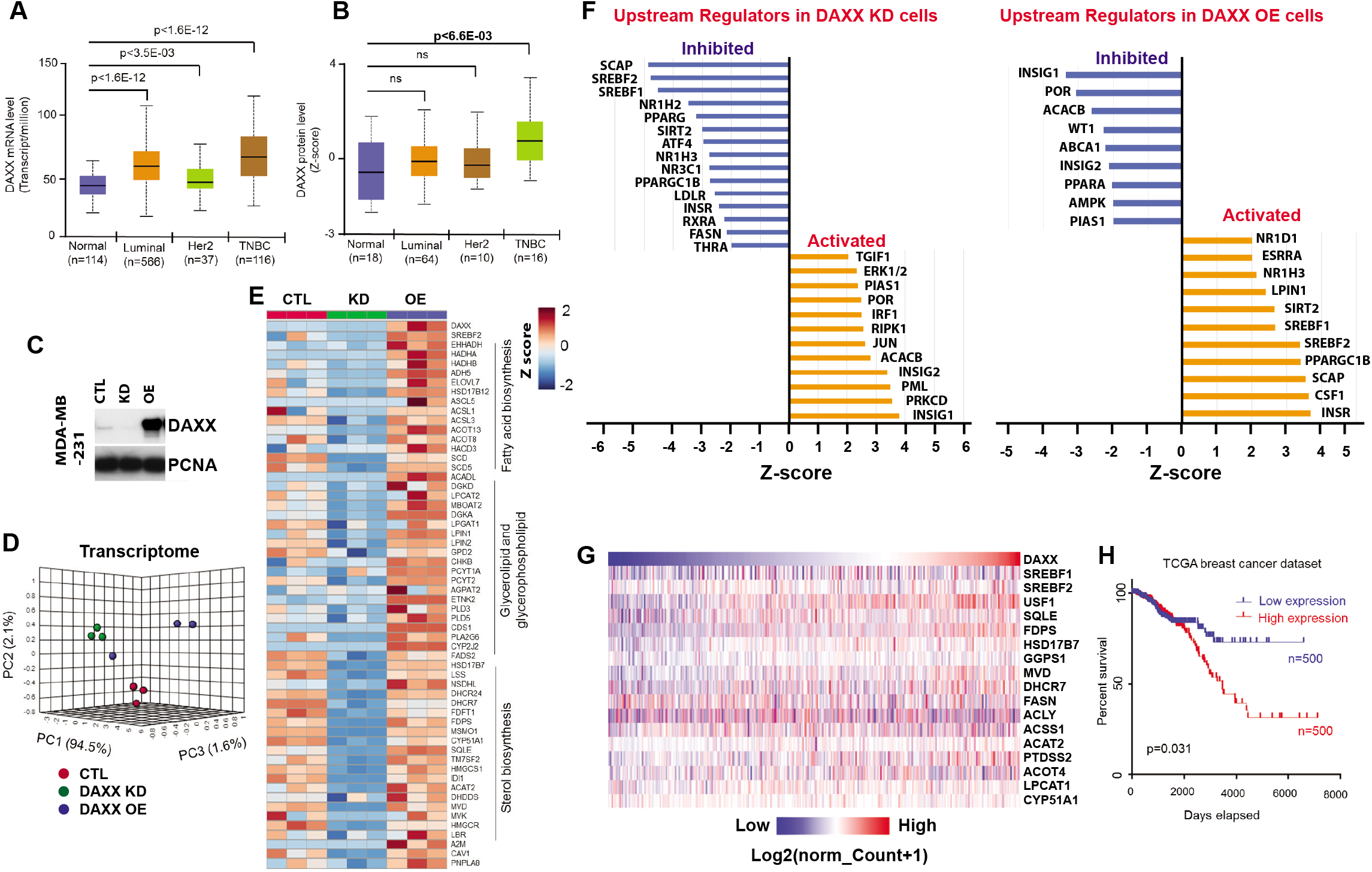
DAXX is upregulated in BC and transcriptomic profiling identifies a functional role for DAXX in lipid metabolism. (**A**) Upregulation of DAXX mRNA expression in three major BC subtypes compared to normal controls based on a TCGA dataset. (**B**) Increased DAXX protein levels in three major BC subtypes compared to normal controls based on a CPTAC dataset. (**C**) Validation of shRNA-mediated DAXX knockdown (KD) and the overexpression of WT DAXX cDNA (OE) compared to cells with a control vector (CTL) in MDA-MB-231 cells by immunoblotting. (**D**) Principal component analysis comparing transcriptomes of CTL, DAXX KD, and WT DAXX OE cells. Each dot represents a sample. (**E**) Hierarchical clustering heatmap analysis of differentially expressed genes showing 30 most differentially expressed genes. (**F**) Top 10 pathways identified using IPA as downregulated in DAXX KD but upregulated in DAXX OE cells. (**G**) A heatmap of relative mRNA levels of DAXX along with select genes in the lipid metabolism pathways. The red and blue groups refer to a high (red) or low level (blue) of mRNA expression of the indicated genes according to combined expression scores in an individual tumor sample from a TCGA BC dataset. (**H**) A Kaplan-Meier plot of the correlation between gene expression levels of the select genes in panel **G** (the red and blue groups) and patient survival time.

RT-qPCR analyses provided validation for the microarray results (Supplementary Figure S2). The impact of DAXX knockdown or overexpression on lipogenic gene expression was further validated by immunofluorescence microscopy and immunoblotting (Supplementary Figure S2B and C). Using a tetracycline-inducible gene expression system, we found that DAXX induction increased lipogenic gene expression (Supplementary Figure S2D), providing further evidence that DAXX directly activates lipogenic gene expression. In keeping with our findings, our analysis of public gene expression datasets based on human and mouse cells (42-44) indicated that DAXX is involved in promoting the SREBP/lipogenesis pathway (Supplementary Figure S3). Further analyses of the TCGA data indicate that DAXX expression levels positively correlate with that of SREBP1 and SREBP2 (Supplementary Figure S5) and a panel of lipogenic genes (Figure 1G). Notably, high mRNA levels of the gene set including DAXX shown in Figure 1G predict poor patient survival (Figure 1H). Collectively, these data provide evidence that DAXX may play an important role in promoting lipogenic gene expression.

### DAXX promotes lipid production

To determine whether transcriptomic differences correspond to an alteration of intracellular lipidome, the same panel of MDA-MB-231-derived cell lines used for our transcriptomic study were subjected to global lipidome analysis using LC-MS technique. Similar to transcriptomic profiles, control, DAXX KD and OE cells cluster into groups with distinct intracellular lipidomes (Figure 2A and B). A lipidome-based pathway analysis again revealed that DAXX expression levels significantly impact lipogenesis pathways (Figure 2C). To validate DAXX’s role in lipogenesis, we depleted DAXX using CRISPR/Cas9 (Supplementary Figure S5A). Lipidomic profiling again showed that DAXX depletion significantly altered lipidomes (Supplementary Figure S5B-E).

**Figure 2.**
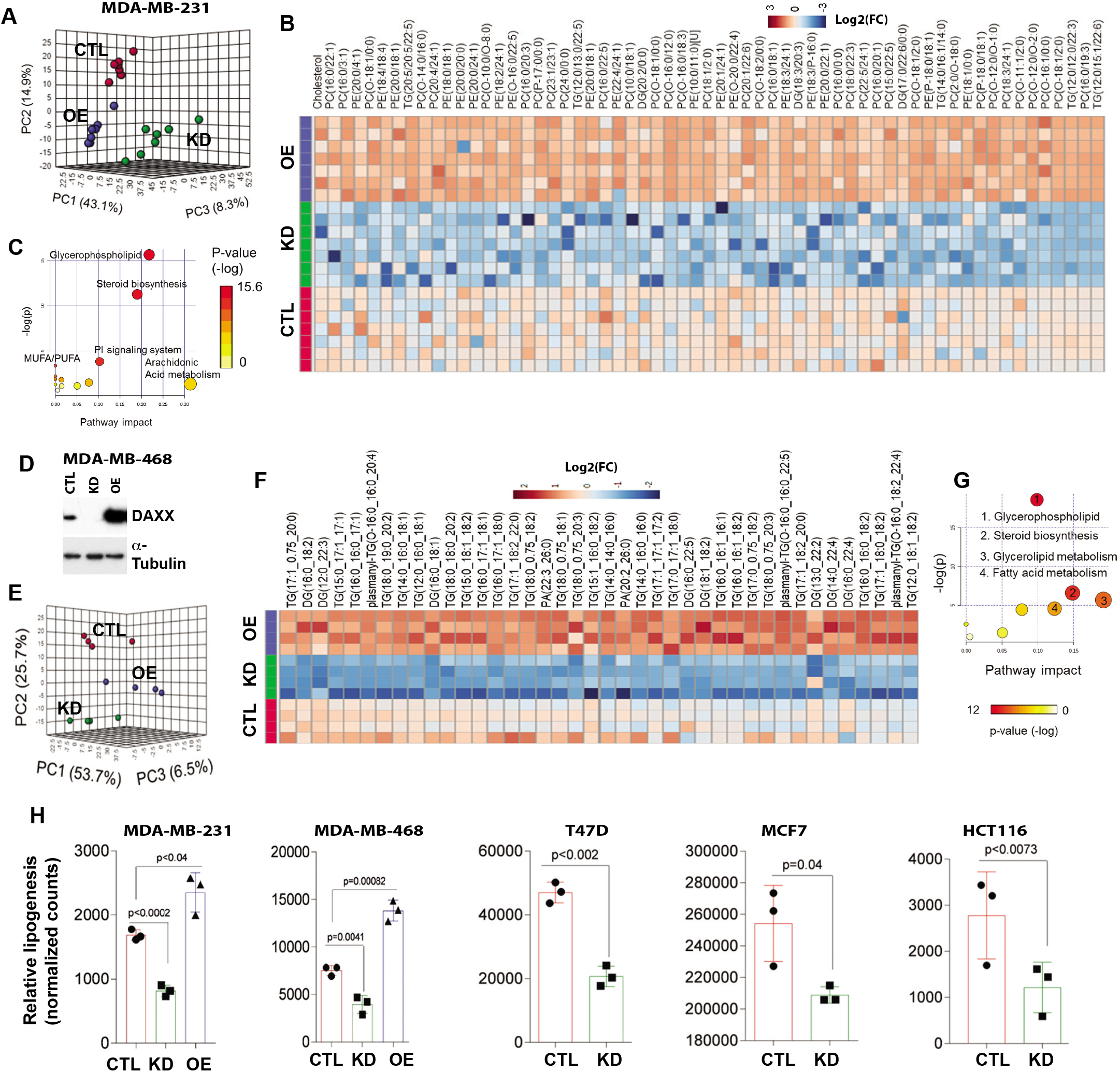
DAXX promotes lipogenesis in cancer cells. (**A**) Principal component analysis comparing lipidomes of MDA-MB-231 cells (CTL, DAXX KD and OE). Each dot represents a sample (n=6). (**B**) Hierarchical clustering heatmap analysis of the 60 most differentially expressed lipid molecules in CTL, KD and OE MDA-MB-231 cells. (**C**) Significantly altered lipid pathways in MDA-MB-231 cells with DAXX OE that were identified using the KEGG pathway library with an FDR <0.05 and a pathway impact >0.5. The color and size of the circle denote p value and pathway impact respectively. The largest red circle indicates the most significantly affected pathway. (**D**) An immunoblotting analysis of MDA-MB-468 cells with a control vector (CTL), DAXX shRNA (KD), and DAXX cDNA (OE). (**E**) Principal component analysis of lipidomes of CTL, KD and OE MDA-MB-468 cells. Each dot represents a sample (n=4). (**F**) Hierarchical clustering heatmap analysis of top differentially changed lipid molecules in MDA-MB-468 cells. (**G**) Significantly altered lipid pathways in MDA-MB-468 DAXX OE cells based on lipidome as in panel **C**. The top 4 most altered pathways are labelled. (**H**) Impact of DAXX expression levels on acetate-dependent de novo lipid synthesis using [^14^C]-acetate labeling in the absence of serum in the indicated cell lines with different levels of DAXX expression (CTL, KD or OE).

To understand the broader role of DAXX in lipogenesis, we have explored correlation of DAXX expression with lipid production in different cancer types. We have depleted endogenous DAXX or overexpressed wild-type DAXX in another TNBC cell line MDA-MB-468 (Figure 2D). Consistent with lipidomic changes in cells derived from MDA-MB-231, global lipidomic profiling revealed that DAXX KD reduced, but WT OE increased levels of diverse lipid molecules, respectively, in MDA-MB-468-derived cells (Figure 2E-G). Principal component analysis (PCA) analysis indicated that MDA-MB-468 cells with different levels of DAXX expression exhibit distinct lipid profiles (Figure 2E). Pathway analyses based on metabolites identified the biosynthesis pathways of glycerophospholipid, steroid, glycerolipid and fatty acid metabolism as the top pathways influenced by DAXX expression levels (Figure 2G).

It is well known that cancer cells utilize acetate as an alternative carbon source to glucose for de novo lipogenesis (6). To assess whether DAXX affects acetate-driven de novo lipogenesis, we treated cells that express different levels of DAXX with [^14^C]-acetate. Quantification of the [^14^C]-labeled lipids showed that DAXX expression levels positively correlated with levels of intracellular lipids, with reduced or increased labeled lipids, respectively, in DAXX KD or DAXX OE in cells derived from MDA-MB-231 and MDA-MB-468 cells (Figure 2H). Diminished de novo lipogenesis upon DAXX depletion was also observed in BC cell lines of luminal subtypes (MCF7 and T47D) and the colon cancer cell line HCT116 (Figure 2H). Collectively, these metabolic experiments established a functional role for DAXX in de novo lipogenesis in cancer cells.

### DAXX is critical for tumor growth and lipogenesis in vivo

As de novo lipogenesis is critical to cell proliferation and tumorigenesis (1,45), DAXX expression levels could impact cell growth in vitro and tumor growth in vivo due to alteration in lipid production. Indeed, DAXX knockdown reduced the number and size of colonies when compared to control, while WT DAXX OE had the opposite effects in three-dimensional cell culture model of MDA-MB-231 (Supplementary Figure. S6).

Next, we examined effects of DAXX expression levels on tumor growth in vivo. In orthotopic BC xenograft models using female mice, DAXX knockdown markedly reduced while WT DAXX OE significantly increased tumor growth of both MDA-MB-231 and MDA-MB-468 TNBC cell lines (Figure 3A and B). Despite the notable difference in the tumor growth rate between the MDA-MB-231 and the MDA-MB-468 xenograft tumors, the effects of DAXX expression levels on tumor growth were clearly observed in both TNBC tumor models (Figure 3A and B). Immunoblotting analysis of tumor extracts showed that DAXX KD and OE were maintained in in vivo (Figure 3C). We profiled the lipids in xenograft tumors derived from cells with different levels of DAXX expression and lipid production. We found that the expression levels of DAXX positively correlated with elevated levels of lipids in xenograft tumors (Figure 3D). PCA analysis indicated that tumors with different DAXX expression levels exhibited distinct lipid profiles (Figure 3E). Overexpression of DAXX significantly elevated glycerophospholipids (Figure 3D), total TGs (n=350), and metabolites involved in cholesterol biosynthesis (Figure 3F). Altogether, our data demonstrate that DAXX promotes oncogenic lipogenesis and tumor growth in vivo.

**Figure 3.**
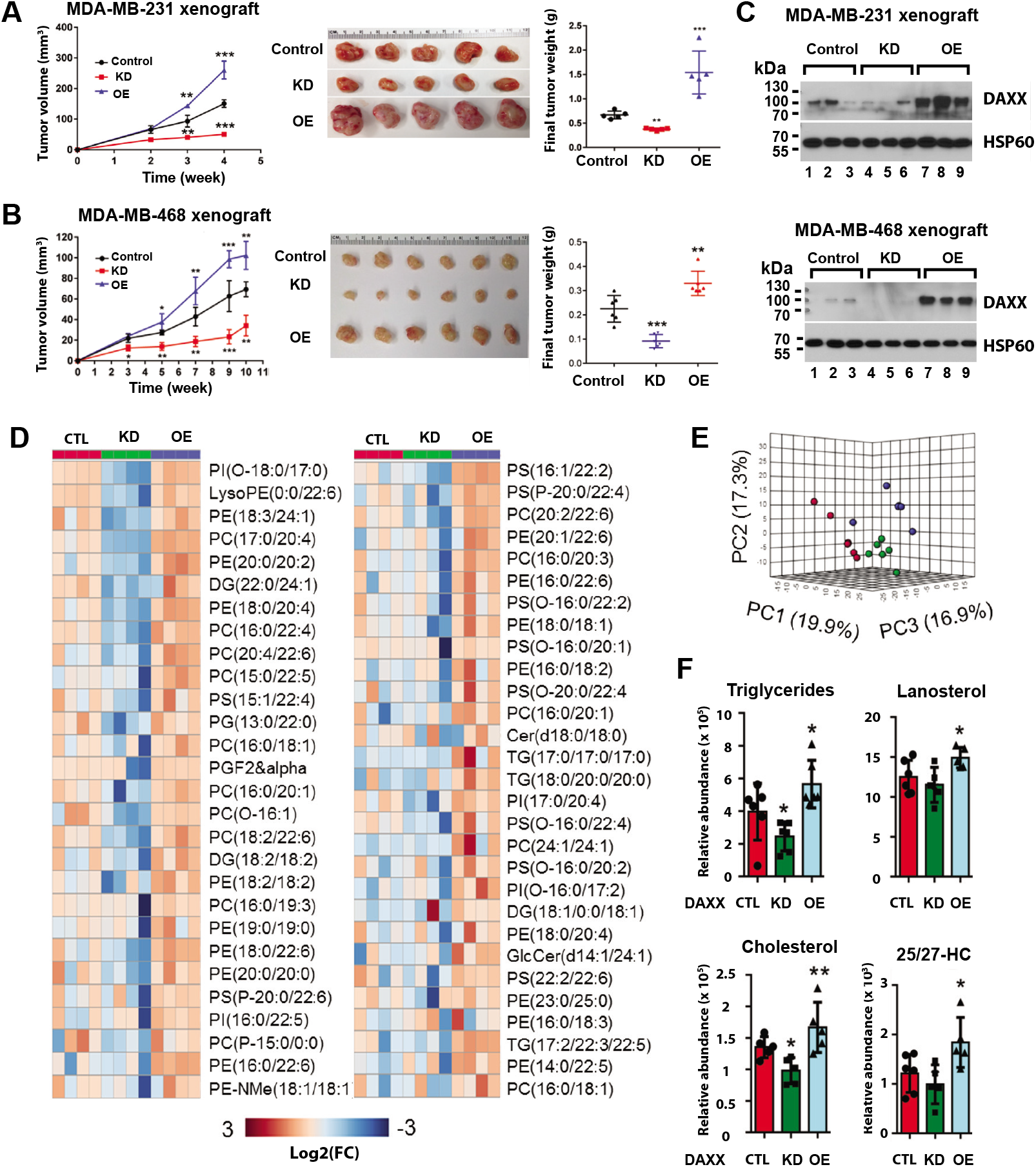
DAXX promotes tumor growth in vivo. (**A** and **B**) Cell lines derived from MDA-MB-231 or MDA-MB-468 stably transduced with a control vector (Control, CTL), DAXX shRNA (KD), or WT DAXX cDNA (OE) were implanted into mammary fat pads of female NSG mice. Longitudinal tumor volumes are plotted. Tumor images and weights at the endpoint are shown. (**C**) DAXX KD and overexpression were maintained in vivo. Protein extracts from three representative xenograft tumors were analyzed for DAXX protein levels using immunoblotting. HSP60 was detected as a loading control. (**D**) Hierarchical clustering heatmap analysis of top glycerophospholipid molecules that were differentially produced in MDA-MB-231 xenograft tumors with different levels of DAXX. (**E**) Multivariate PCA of lipids shows distinct global lipid profiles in xenograft tumors derived from control (red dots), DAXX KD (green dots), and OE (blue dots) MDA-MB-231 cells. (**F**) Relative abundance of total triglycerides, cholesterol and derivatives in xenograft tumors derived from MDA-MB-231 cell line panel as in (**A**). Box plots of the indicated lipid species are shown. The p values were calculated based on Student’s t-test. *: p < 0.05; **: p < 0.01. 25/27-HC: 25- or 27-hydroxycholesterol.

### DAXX interacts with SREBP1 and SREBP2

SREBP1/2 are master transcription factors that promote lipid production when the intracellular levels of lipids/sterols are low (3,4,46). Because DAXX expression levels positively correlate with the activation of SREBP/lipid biosynthesis pathway (Figures 1–3), we reasoned that DAXX could regulate lipid biosynthesis through interacting with SREBPs. Immunoprecipitation (IP) of total cell extracts with an anti-DAXX antibody evidenced co-precipitation of the precursor and mature (M) forms of SREBP2 (Figure 4A). Using Proximity Ligation Assay (PLA) with a mouse monoclonal anti-DAXX and a rabbit polyclonal anti-SREBP2 antibody, endogenous DAXX-SREBP2 interaction signals were detected in both the cytoplasm and the nucleus of the MDA-MB-231 cells (Figure. 4B). Interestingly, the number of DAXX/SREBP2 PLA signals was significantly increased in the absence of serum (Figure 4B). The number of PLA signal dots were shown to be proportional to cellular protein levels (31). Thus, low extracellular supply of lipids appeared to enhance the DAXX-SREBP2 interaction.

**Figure 4.**
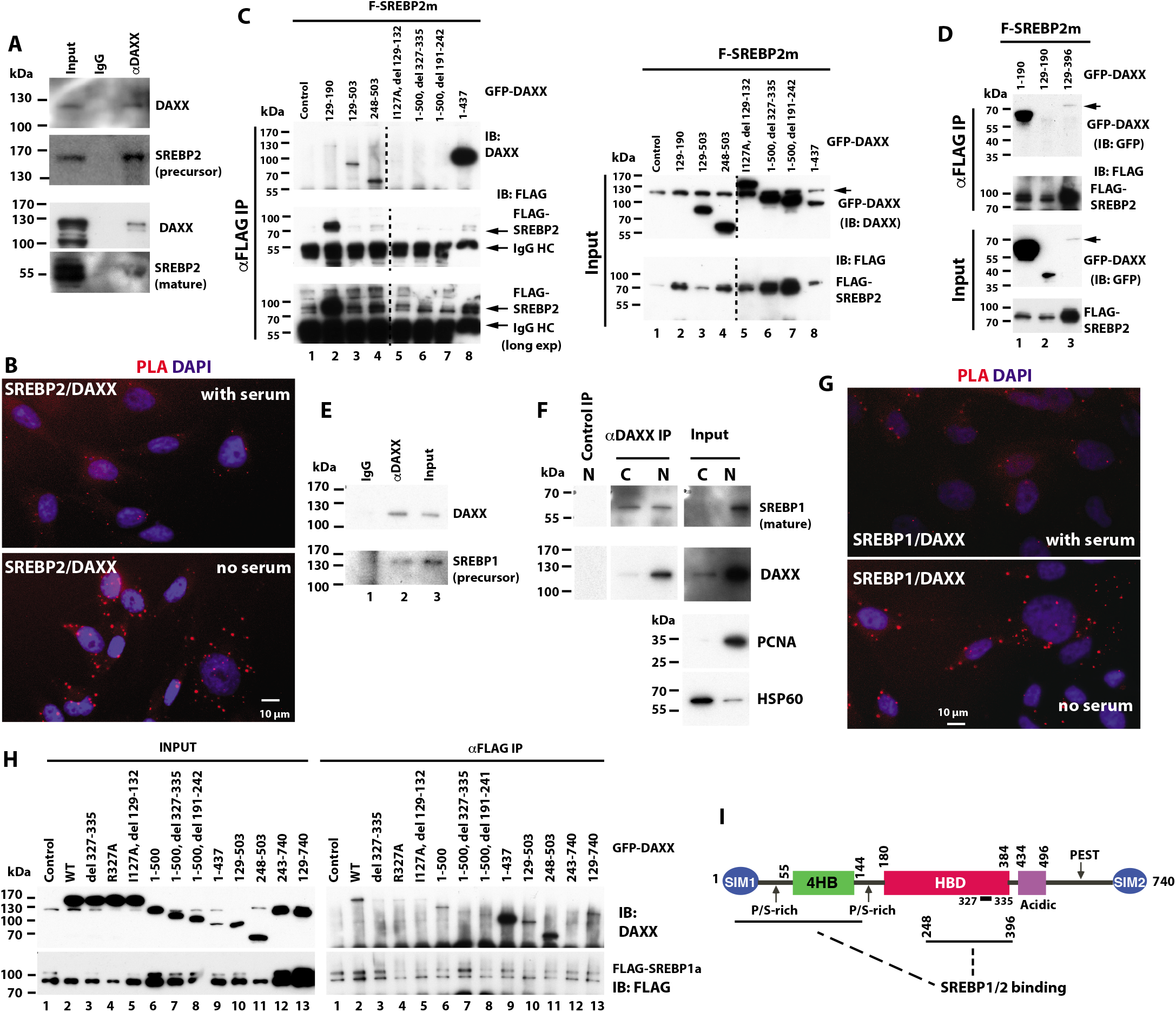
DAXX binds to SREBPs. (**A**) The endogenous DAXX and SREBP2 interact. MDA-MB-231 whole cell extracts were subjected to IP with a control (IgG) or an anti-DAXX antibody. The immunoprecipitated SREBP2 and DAXX were detected. (**B**) Representative images of Proximity Ligation Assay (PLA) showing DAXX-SREBP2 interactions in MDA-MB-231 cells in the presence or absence of serum. (**C** and **D**) There are two independent binding sites in DAXX for mature SREBP2. 293T cells were cotransfected with FLAG-SREBP2m (mature) and GFP (control) or an indicated GFP-DAXX fusion construct. The cell lysates were subjected to anti-FLAG IP. Note that the DAXX amino acid (aa) 129-190 construct is not recognized by the antibody used for detecting DAXX in **C** (lane 2), which was detected with a GFP antibody (panel **D**, lane 2). The endogenous DAXX in the input samples is denoted with an arrow in **C**. In **D**, the arrow points to the GFP-DAXX 129-396 band, which accumulated at a relatively low level. HC: heavy chain. (**E** and **F**) The endogenous DAXX and SREBP1 interact. Total MDA-MB-231 cell extracts (**E**), the cytoplasmic (C) or nuclear fraction (N) were subjected to IP was in panel **A** and the co-precipitated SREBP1 and DAXX were detected. PCNA and HSP60 were detected as a marker of nuclear and cytoplasmic fraction respectively in panel **F**. (**G**) PLA images showing DAXX-SREBP1 interactions in MDA-MB-231 cells in the presence or absence of serum as in panel **B**. (**H**) Cotransfection of FLAG-SREBP1a (mature) and the indicated GFP-DAXX fusion constructs, IP and immunoblotting experiments were performed as in **C**. (**I**) Schematic drawing of DAXX-SREBP interactions. The position of aa 327-335 within the DAXX HBD critical for the DAXX-SREBP interactions is indicated. SIM: SUMO-interacting motif; 4HB: DAXX helical bundle; HBD: histone-binding domain; PEST: proline, glutamic acid, serine, and threonine-rich sequence. Numbers refer to aa residue positions in the DAXX protein. In panel **C**, vertically sliced images from the same gel are juxtaposed as indicated.

Likewise, DAXX interacted with both the precursor and mature forms of SREBP1 in MDA-MB-231 cells (Figure 4E and F). To assess the interaction between DAXX and mature SREBP1, we conducted co-IP using nuclear and cytoplasmic fractions of the MDA-MB-231 cells. As expected, the mature SREBP1 was predominantly detected in the nucleus (Figure 4F). Notably, mature SREBP1 was enriched in the DAXX immunoprecipitates of both fractions (Figure 4F). The DAXX-SREBP1 interaction signals were also detected in MDA-MB-231 cells using PLA (Figure 4G). Similar to the DAXX-SREBP2 interaction (Figure 4B), the number of DAXX-SREBP1 PLA foci was increased in the absence of serum (Figure 4G). Altogether, these data show that DAXX binds to both precursor and mature SREBP1/2.

Using various DAXX deletion constructs in transfected 293T cells, we found that the mature SREBP2 interacted with two separate regions of DAXX, the N-terminal part encompassing the well-folded helical bundle domain termed 4HB (DAXX helical bundle) (47) and a part of the central histone-binding domain (HBD) (24) (Figure 4C and D). Interestingly, although 4HB and HBD individually bound robustly to SREBP2 (Figure 4C lanes 3 and 4; Figure 4D lanes 1 and 3), the integrity of both binding sites in the context of the full-length DAXX or a longer construct appeared critical for the DAXX-mature SREBP2 interaction. Indeed, mutations within 4HB (I127A, del 129-132) (Figure 4C, lane 5) or HBD (del 327-335, Figure 4C lane 6; del 191-242, Figure 4C lane 7) abolished the DAXX-SREBP2 interaction. Notably, the DAXX construct (aa 1-437) lacking the sequence from the acidic domain to the C-terminus seemed to show higher affinity to SERBP2 (Figure 4C lane 8). As deletions of C-terminal regions of DAXX did not affect the DAXX-SREBP2 interaction (Figure 4C and D) and C-terminal fragments spanning aa 574-740 did not bind to SREBP2 (data not shown), we concluded that SREBP2 does not bind to the C-terminal domain of DAXX. The mature SREBP1a bound to DAXX in a similar fashion (Figure 4H). Collectively, our data demonstrate that the mature SREBP1/2 specifically interact with DAXX via DAXX’s 4HB and HBD (Figure 4I).

### SREBP-binding sites are enriched in DAXX-associated chromatins

The data presented above suggest that DAXX promotes SREBP-mediated transcription to stimulate lipogenesis. To test this idea, we conducted luciferase reporter assays. As shown in Figure 5A, forced expression of mature SREBP2, SREBP1a and SREBP1c increased the activity of the luciferase reporter that is under the control of the *SREBF2* promoter containing a canonical *SRE*. Co-expression of DAXX further increased the luciferase activity, while DAXX alone had only minimal effects (Figure 5A).

**Figure 5.**
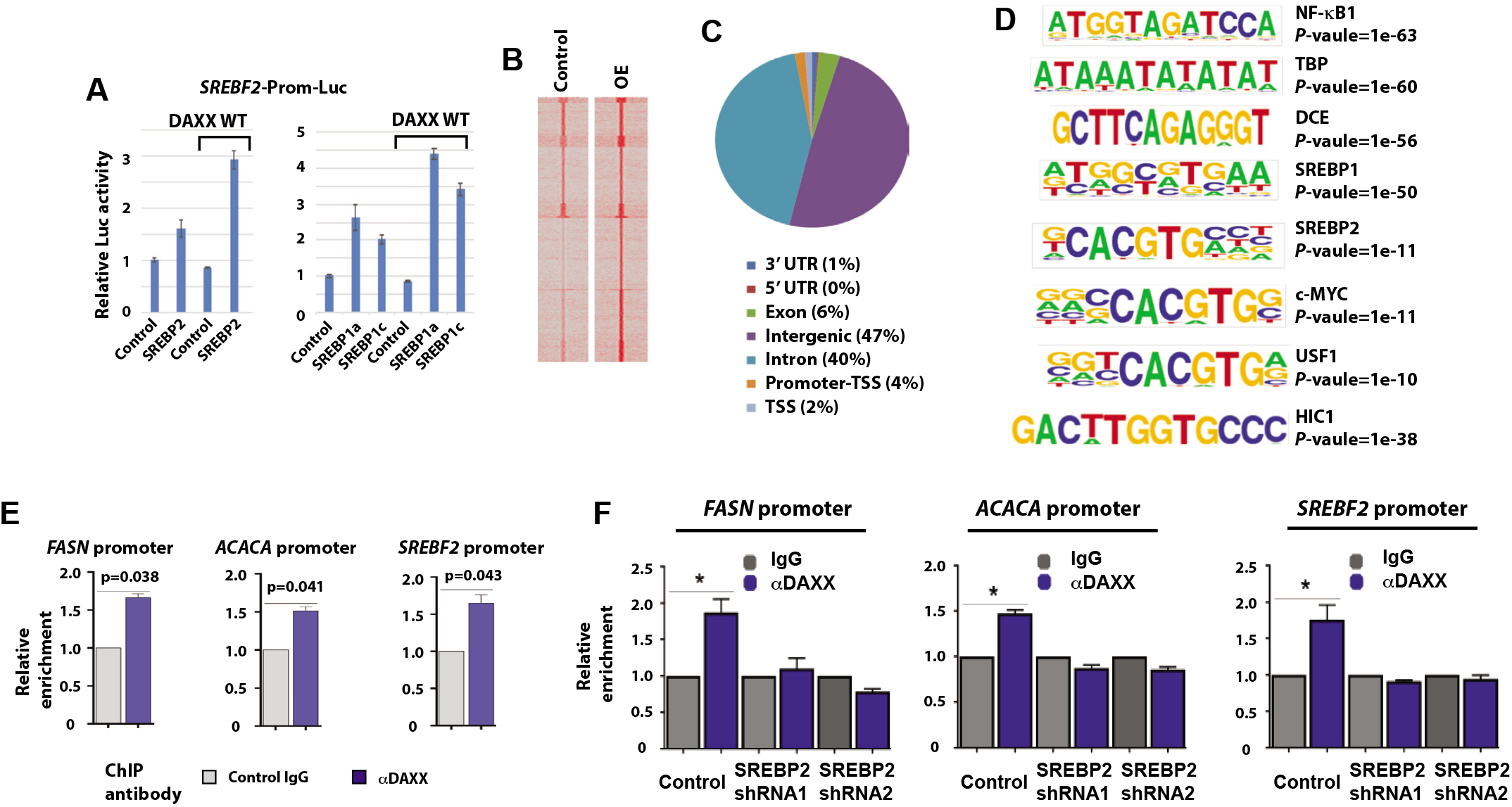
DAXX activates SREBP-mediated transcription and occupies the promoters of lipogenic genes. (**A**) MDA-MB-231 cells were transfected with a luciferase reporter driven by a promoter fragment from the *SREBF2* gene along with mature SREBP2, SREBP1a, SREBP1c, or wt DAXX cDNA as indicated. Dual luciferase assays were done. (**B**) ChIP-seq signal intensity heat maps in MDA-MB-231 control and DAXX OE cell lines; signals are centralized to transcriptional start sites (TSS). (**C**) The genome-wide distribution of DAXX chromatin occupancy. (**D**) Motifs enriched as determined by the DAXX ChIP-seq dataset of MDA-MB-231 DAXX OE cells. (**E**) MDA-MB-231 cells stably expressing WT DAXX were subjected to ChIP with a control IgG and an anti-DAXX antibody. The precipitated DNAs were subjected to qPCR with primers specific to promoter regions of the indicated genes. (**F**) SREBP2 is critical for DAXX to bind lipogenic gene promoters. MDA-MB-231 cells with a control vector or a SREBP2 shRNA vector were subjected to ChIP with a control IgG, or an anti-DAXX antibody followed by qPCR with primers specific to the indicated gene promoters.

We surveyed genome-wide occupancy of DAXX using the ChIP-seq technology. Overexpression of WT DAXX increased DAXX’s chromatin association (Figure 5B). Consistent with other studies (42), DAXX primarily bound to sites in introns and intergenic regions with less frequent association with promoters (Figure 5C). A de novo motif analysis revealed that SREBP-binding elements were significantly enriched in DAXX-associated sites (Figure 5D and Supplementary Figure S7).

ChIP-qPCR experiments demonstrated that WT DAXX was enriched in the promoters of *FASN, ACACA* and *SREBF2* (Figure 5E). In MDA-MB-231 cells depleted of SREBP2, DAXX’s recruitment to the promoters of *FASN, ACACA* and *SREBP2* was impaired (Figure 5F), demonstrating that SREBP2 is critical for DAXX to bind the promoters of lipogenic genes. Likewise, DAXX recruitment to the promoters of lipogenesis genes were impaired in MDA-MB-231 cells depleted of SREBP1 (Data not shown). Altogether, these data demonstrating that SREBPs are critical for DAXX to bind the promoters of lipogenic genes.

### The DAXX-SREBP axis is important for lipogenesis and tumor growth

SREBP1/2 drive lipid biosynthesis to promote tumorigenesis (4,46). In MDA-MB-231 cells, SREBP2 knockdown reduced de novo lipogenesis from acetate and tumor growth in vivo (Figure 6A and E), whereas the overexpression of mature SREBP2 increased lipogenesis and tumor growth (Figure 6A and E). Concordantly, lipidomic profiling shows that SREBP2 knockdown had a marked impact on global lipid landscapes (Figure 6B-D). Likewise, SREBP1 knockdown (or overexpression) impaired (or promoted) de novo lipogenesis and tumor growth, respectively (data not shown). SREBP1 knockdown also significantly altered intracellular lipidome (Supplementary Figure 8). These data indicate both SREBP1 and SREBP2 are critical mediators of lipogenesis and tumorigenesis in TNBC models. Our data presented above demonstrated that the DAXX-SREBP interactions are critical for lipogenic gene expression, de novo lipogenesis and tumor growth. To further link SREBP2 to DAXX-mediated tumorigenesis, we depleted SREBP2 in MDA-MB-231 cells with WT DAXX OE. We observed that SREBP2 knockdown in the DAXX OE cells significantly attenuated the levels of lipogenic enzymes and tumor growth (Figure 6F and G), suggesting that SREBP2 is a critical effector of DAXX’s oncogenic function.

**Figure 6.**
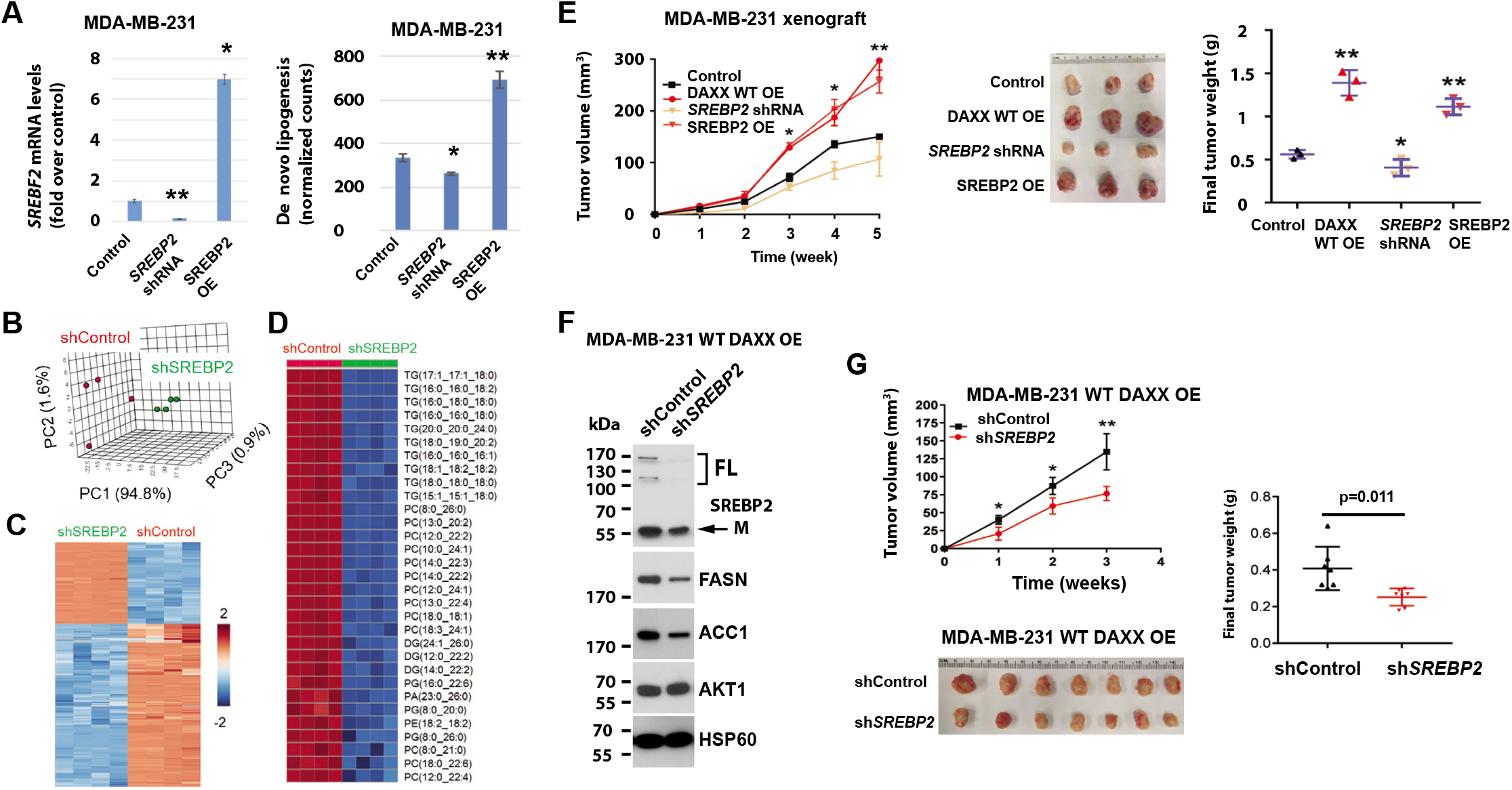
SREBP2 knockdown impairs DAXX-mediated tumor growth. (**A**) Cells derived from MDA-MB-231 cell line with a vector for a control, an SREBF2 shRNA, or SREBP2 (mature) cDNA were subjected to RT-qPCR for assessing SREBP2 expression, and de novo lipogenesis assays using [^14^C] acetate. (**B**) PCA of lipidomes in shControl and shSREBP2 cells. Each dot represents a sample (n=4). (**C**) Hierarchical clustering heatmap analysis demonstrates global lipid landscape in shSREBP2 cells compared to shControl cells. (**D**) Hierarchical clustering heatmap analysis of lipids including glycerolipid and glycerophospholipid molecules that were highly downregulated in shSREBP2 cells compared to control cells. (**E**) MDA-MB-231-derived cells (Control, DAXX OE, SREBP2 shRNA and mature SREBP2 OE) were xenografted into mammary fat pads of female NSG mice. Tumor volumes were plotted against time. Representative images of dissected tumors are shown. The final tumor weights are plotted. (**F**) Control or SREBF2 shRNA were expressed in MDA-MB-231 cells with DAXX OE. The levels of the indicated proteins were assessed by immunoblotting. (**G**) The indicated cells shown in panel **F** were xenografted into mammary fat pads of female NSG mice. Tumor volumes were plotted against time. Representative images of dissected tumors are shown. The final tumor weights are plotted. The p values were calculated (vs. control) based on Student’s t-test. *: p < 0.05; **: p < 0.01.

To further assess the importance of the DAXX-SREBP interaction on lipogenesis, we overexpressed a DAXX mutant (del 327-335) defective of SREBP1 or SREBP2 binding (Figure 4C and H) in MDA-MB-231 cells. The protein levels of both WT DAXX and the del 327-335 mutant were similar (Figure 7A and B). A de novo lipogenesis assay using [^14^C]-acetate labeling indicated that the del 327-331 mutant attenuated lipogenesis (Figure 7C). Lipidomic profiling revealed that MDA-MB-231 cells expressing the del 327-335 mutant has a distinct global lipid profile from that of cells expressing the WT DAXX (Figure 7D and E) and that this mutant was impaired to enhance lipid production including glycerolipids and glycerophospholipids as compared to WT DAXX (Figure 7F and G). Reduced levels of specific lipid molecules such as cholesterol and fatty acid derivatives were evident in MDA-MB-231 cells expressing the del 327-331 mutant compared to those expressing the WT DAXX (Figure 7H). In vivo, the growth of xenograft tumors derived from cells expressing the del 327-335 mutant was markedly slower than that derived from cells with WT DAXX (Figure 7I). These data collectively indicate that the DAXX-SREBP interaction is critical for DAXX to promote lipogenesis and tumorigenesis (Figure 7J).

**Figure 7.**
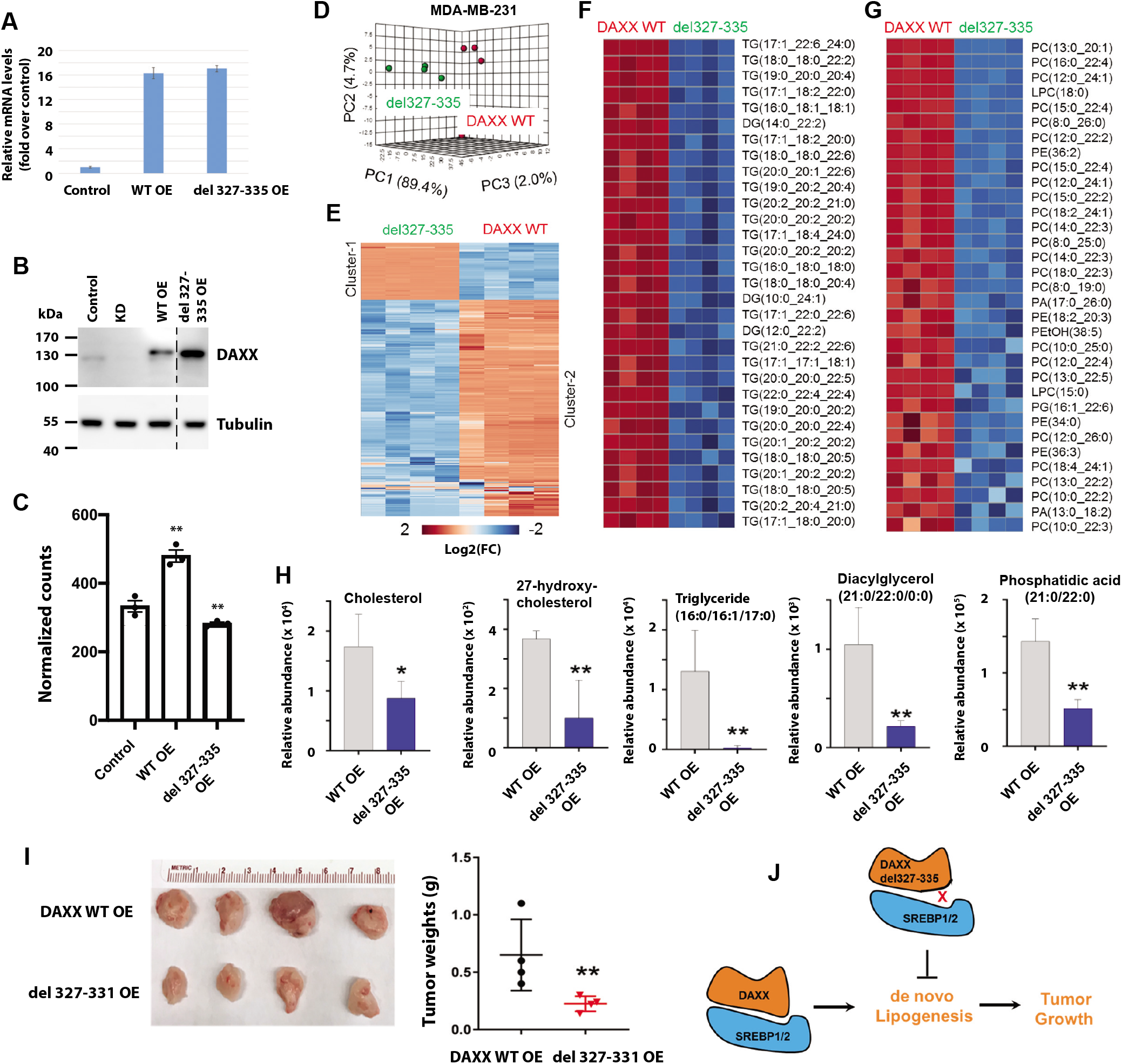
The DAXX-SREBP interaction is critical for lipogenesis and tumor growth. (**A**) Relative mRNA levels of DAXX in MDA-MB-231 cells expressing the WT or del 327-335 mutant cDNA of DAXX as determined by RT-qPCR. (**B**) Protein levels of DAXX in control cells and those with DAXX KD, WT and del 327-335 mutant cDNA of DAXX. (**C**) The DAXX del 327-337 mutant impaired de novo lipogenesis. Serum-starved cells were labeled with [^14^C] acetate and total lipids were isolated. Radioactivity was counted and normalized against total protein level. (**D**) PCA of lipidomes in MDA-MB-231 cells expressing the del327-335 mutant and WT DAXX. Each dot represents a sample (n=4). (**E**) Hierarchical clustering heatmap analysis of global lipidomes in cells expressing the del327-335 mutant and WT DAXX. (**F**) Hierarchical clustering heatmap analysis of glycerolipid molecules that were highly differentially expressed between MDA-MB-231 cells with the del 327-335 mutant and wt DAXX. (**G**) Hierarchical clustering heatmap analysis of glycerophospholipid molecules that were highly differentially expressed between MDA-MB-231 cells with the del 327-335 mutant and WT DAXX. (**H**) Bar graphs of relative normalized abundance of specific lipids in MDA-MB-231 cells expressing the del 327-335 mutant and WT DAXX. (**I**) MDA-MB-231 cells expressing the del 327-335 mutant and WT DAXX were xenografted into mammary fat pads of female NSG mice. Representative images of dissected tumors are shown. The final tumor weights are plotted. (**J**) A cartoon depicting the importance of DAXX-SREBP interaction for lipogenesis and tumorigenesis. The p values were calculated (vs. control) based on Student’s t-test. *: p < 0.05; **: p < 0.01.

## DISCUSSION

Lipid availability for proliferating cells determines activity of intracellular lipid biosynthesis pathway. In a nutrient-poor tumor microenvironment, limited supplies of lipids necessitate the activation of intracellular lipid production in tumor cells for sustained tumor growth. An elaborate sterol sensing machinery controls nuclear translocation of SREBP1/2, which promote expression of enzymes required for de novo lipogenesis (1,48). SREBP1/2 in conjunction with other transcription factors, such as the *E*-box-binding basic helix-loop-helix (bHLH) transcription factor USF1, activate expression of lipogenic enzymes and regulators (41). Other coregulators of gene expression such as acetyltransferases (e.g., p300 and PCAF) as well as oncogenic signaling pathways (e.g., KRAS and mTOR) also play important roles in stimulating de novo lipogenesis (6,49). We demonstrated here that DAXX is important for de novo lipogenesis. Mechanistically, DAXX interacts with SREBP1/2 and is enriched in chromatins containing *SRE* motifs. Importantly, the DAXX del 327-335 mutant that cannot bind SREBP1/2 were unable to promote lipogenesis and tumor growth. SREBP2 downregulation prevents enhanced tumor growth by DAXX overexpression. Thus, it is likely that DAXX enhances lipogenesis through interacting with SREBP1/2 to promote lipogenic gene expression, lipid production and tumorigenesis.

Our data suggest that DAXX acts to promote SREBP-mediated transcription. It has been well documented that DAXX can activate and repress transcription, depending on co-regulators that are associated with DAXX (18). Epigenetic modifiers such as HDACs and DNA and histone methyltransferases are involved in DAXX-mediated transcription repression, while coactivators (e.g., CBP) are involved in DAXX-mediated gene activation. The H3.3 histone chaperone function of DAXX is also implicated in both transcriptional activation (50,51) and repression (52,53). Independently of H3.3 deposition by DAXX to chromatin, the H3.3/H4 dimer metabolically stabilizes DAXX protein, which indirectly enhances repression of endogenous retroviruses by a complex consisting of the DAXX-H3.3/H4 sub-complex, HDAC1, KAP1, and SETDB1 (44). Our data show that SREBP1/2 bind to DAXX by contacting with both 4HB and HBD (Figure 4). A previous structural study demonstrates that a peptide within the transactivation domain of p53 binds to DAXX 4HB (47). It will be interesting to assess whether DAXX also engages the transactivation domain of SREBPs to promote transcription and whether the H3.3 chaperone function of DAXX is important for lipogenic gene expression.

Of note, the binding motifs of other known DAXX-binding transcription factors such as NF-κB (20) were highly enriched in DAXX ChIP-seq peaks (Figure 5D). Our ChIP-seq data also implicate the chromatin recruitment of DAXX by other transcription factors such as RUNX1, RUNX2, HIC1 and c-MYC that were not previously shown to interact with DAXX. Furthermore, DAXX might interact with the core-transcriptional machinery, as the TATA-box and DCE (downstream core element) were enriched in DAXX-binding chromatins (Figure 5D and Supplementary Figure S7). These observations suggest a broader role for DAXX in transcription regulation.

Based on our co-immunoprecipitation and PLA results, DAXX appears to interact with SREBP1/2 in both the cytoplasm and the nucleus (Figure 4). Interestingly, the number of DAXX-SREBP1/2 PLA complexes increases upon serum starvation (Figure 4), suggesting that a low level of lipid supply might trigger the formation of the DAXX-SREBP1/2 complexes in the cytoplasm. In the nucleus, the DAXX-SREBP interactions are expected to mediate DAXX’s chromatin recruitment and the activation of lipogenic gene expression. The functional effects of DAXX-SREBP interactions in the cytoplasm are currently unknown. In the cytoplasm, DAXX has been shown to interact with regulators of cell death and cell survival (18). A recent study demonstrates that DAXX promotes the formation of SQSTM1/p62 membrane-less liquid compartments to activate cellular anti-oxidative stress response (54). The functional ramification of the interaction between DAXX and SREBPs in the cytoplasm requires further investigation.

Targeting the de novo lipogenesis pathways such as the biosynthesis of fatty acids (1) and cholesterol (55,56) is a promising approach for treating BC and other cancer types. Our data show that the DAXX-SREBP axis appears to play a critical role in promoting lipid production and tumor growth. The specific binding interactions between DAXX and SREBP1/2 could be potential therapeutic targets for anticancer drug development. It will be interesting to define precisely the interfaces between DAXX and SREBPs and to determine whether these interfaces are tractable as therapeutic targets.

## ACKNOWLEDGEMENTS

We thank Yue Li for help with initial microarray data analysis, Maria Zajac-Kaye, Shuang Huang and Scott Dehm for providing reagents. We also thank Subramaniam Shyamalagovindarajan and Ranjan Perera for library construction and high throughput sequencing for the ChIP-seq experiments.

## Author contributions

I.M. designed, and performed experiments, analyzed and interpreted data, and contributed to writing; G.T., J.W., J.L., A.W., M.L.L., L.Y.Z., and H.T.P. conducted experiments, acquired and analyzed data; J.J.L. performed IPA analysis; T.G. supervised mass spectrometry experiments; Z.H. conducted bioinformatics analysis. Y.D. analyzed and interpreted data and contributed to writing; D.L. designed and conducted experiments, analyzed and interpreted data, supervised the entire study, and wrote the paper.

## Mahmud et al. Supplementary Figures and Tables

**Supplementary Figure S1.**
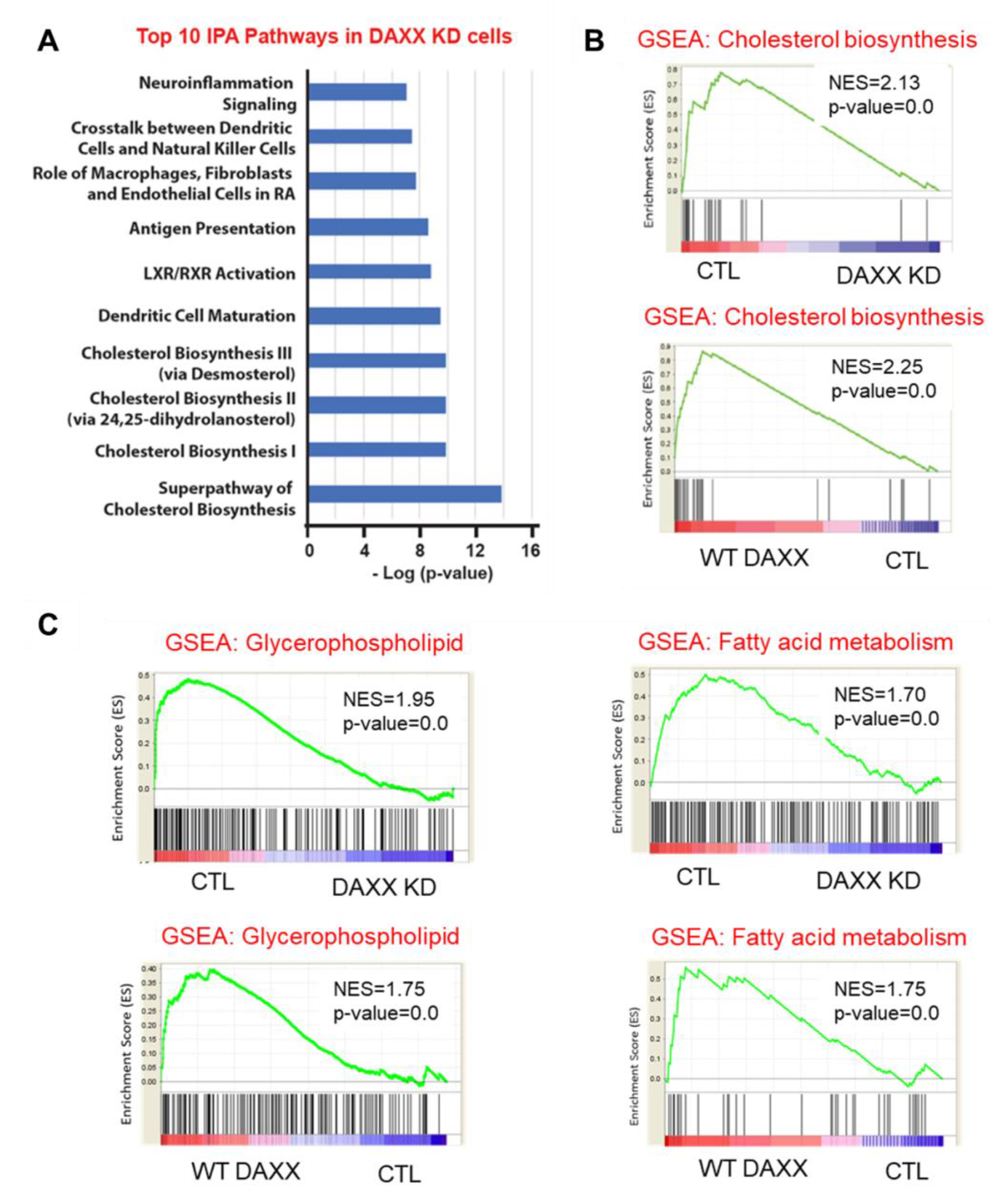
DAXX expression correlates with de novo lipogenesis pathway. (A) Ingenuity pathway analysis (IPA) using differentially expressed genes in DAXX KD cells compared to CTL cells identifies de novo lipogenesis as the most perturbed canonical pathways. (B) Gene set enrichment analyses (GSEA) shows downregulation or upregulation of genes in the cholesterol biosynthesis in MDA-MB-231 cells with DAXX KD or WT DAXX OE, respectively. The KEGG and Reactome genesets were used for the GSEA plots. (C) Gene set enrichment analyses (GSEA) shows downregulation or upregulation of genes in the fatty acid, glycerophospholipid, and glycerolipid metabolism in MDA-MB-231 cells with DAXX KD or WT DAXX OE, respectively. The KEGG and Reactome genesets were used for the GSEA plots.

**Supplementary Figure S2.**
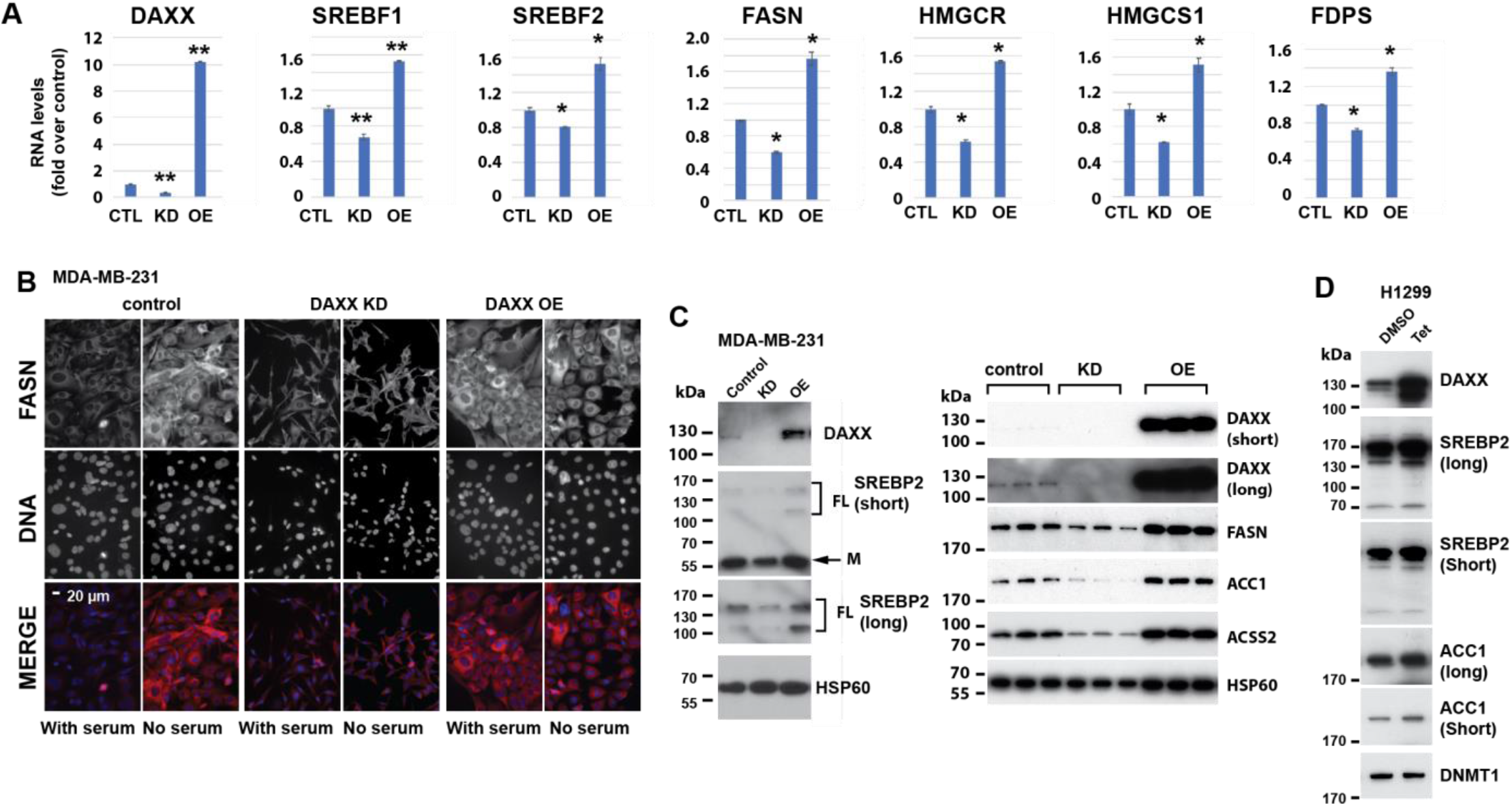
DAXX promotes lipogenic gene expression. (A) Total RNAs were isolated from MDA-MB-231 cells stably transfected with a control vector (CTL), a DAXX shRNA (KD), the WT DAXX cDNA (OE) and subjected to RT-qPCR analysis. The mRNA levels of the indicated genes were normalized against that of ACTB. Data are shown as mean of fold-changes vs. control (CTL) ± SEM (n=3). *: p< 0.05, **: p<0.01 (t-test vs CTL). (B) MDA-MB-231-derived cells were cultured in the presence of serum or serum-starved for 24 hours. The cells were then fixed and stained with an anti-FASN polyclonal antibody (red) and counterstained with DAPI for visualizing nuclei (blue). The cells were imaged using a fluorescence microscope. All images were captured with the same duration of light exposure for the red or blue channel. (C) Immunoblotting analysis of cell extracts of the MDA-MB-231-derived cell lines with antibodies against the indicated proteins. (D) Immunoblotting analysis of cell extracts of the H1299 cells expressing tetracycline (Tet)-inducible wt DAXX in the presence of control (DMSO) or Tet with antibodies against the indicated proteins.

**Supplementary Figure S3.**
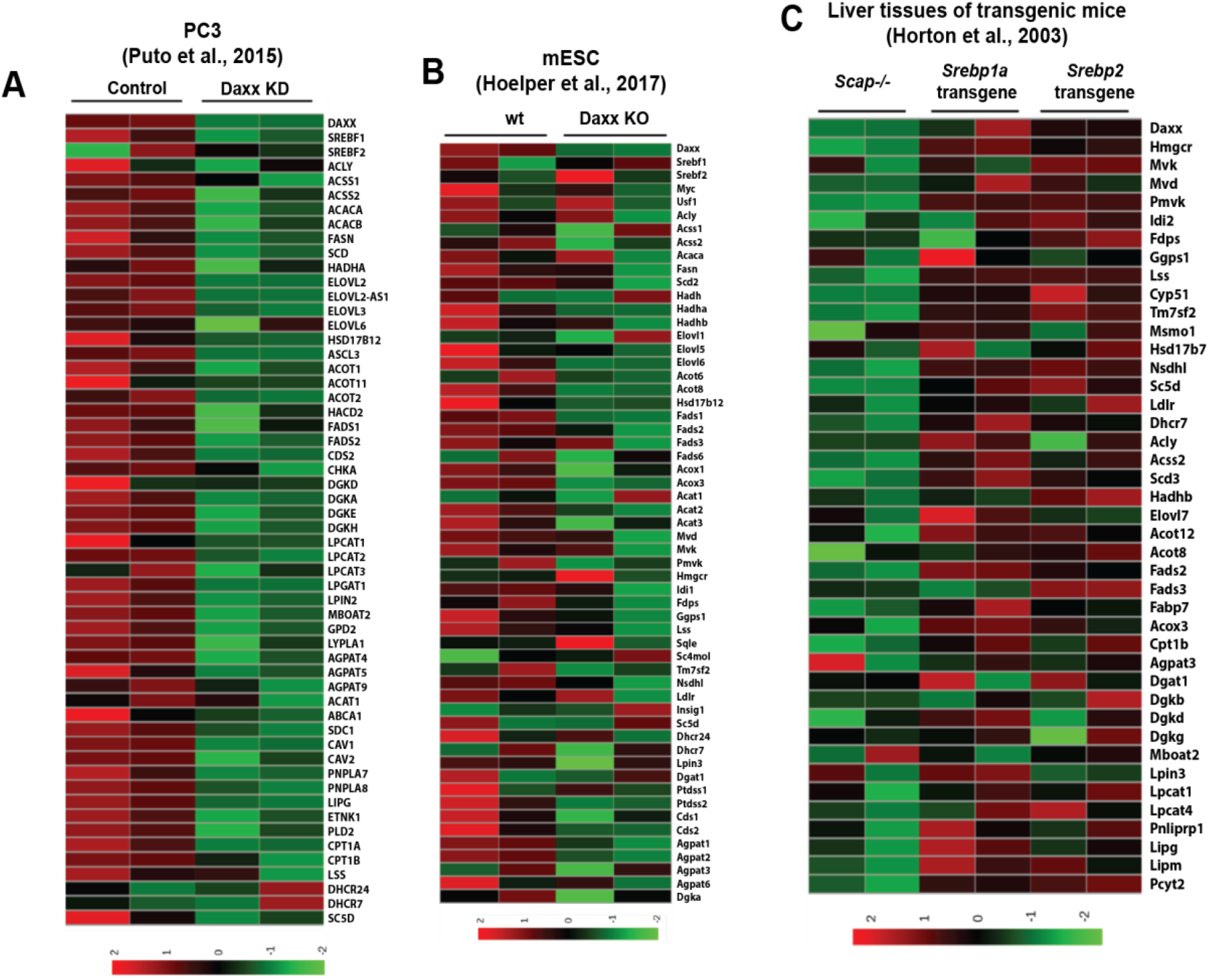
DAXX, SREBP1 and SREBP2 are key regulators for lipogenic gene expression. **(A)** Heatmap of the indicated lipogenic genes in control and DAXX KD cells of the human prostate cancer PC3 cell line. **(B)** Heatmap of the indicated lipogenic genes in wt and Daxx KO cells of the mouse embryonic stem cells (mESC). **(C)** Heatmap of the indicated lipogenic genes in liver tissues from mice with Scap deletion (Scap-/-), the nuclear form of Srebp1a or Srebp2 transgene.

**Supplementary Figure S4.**
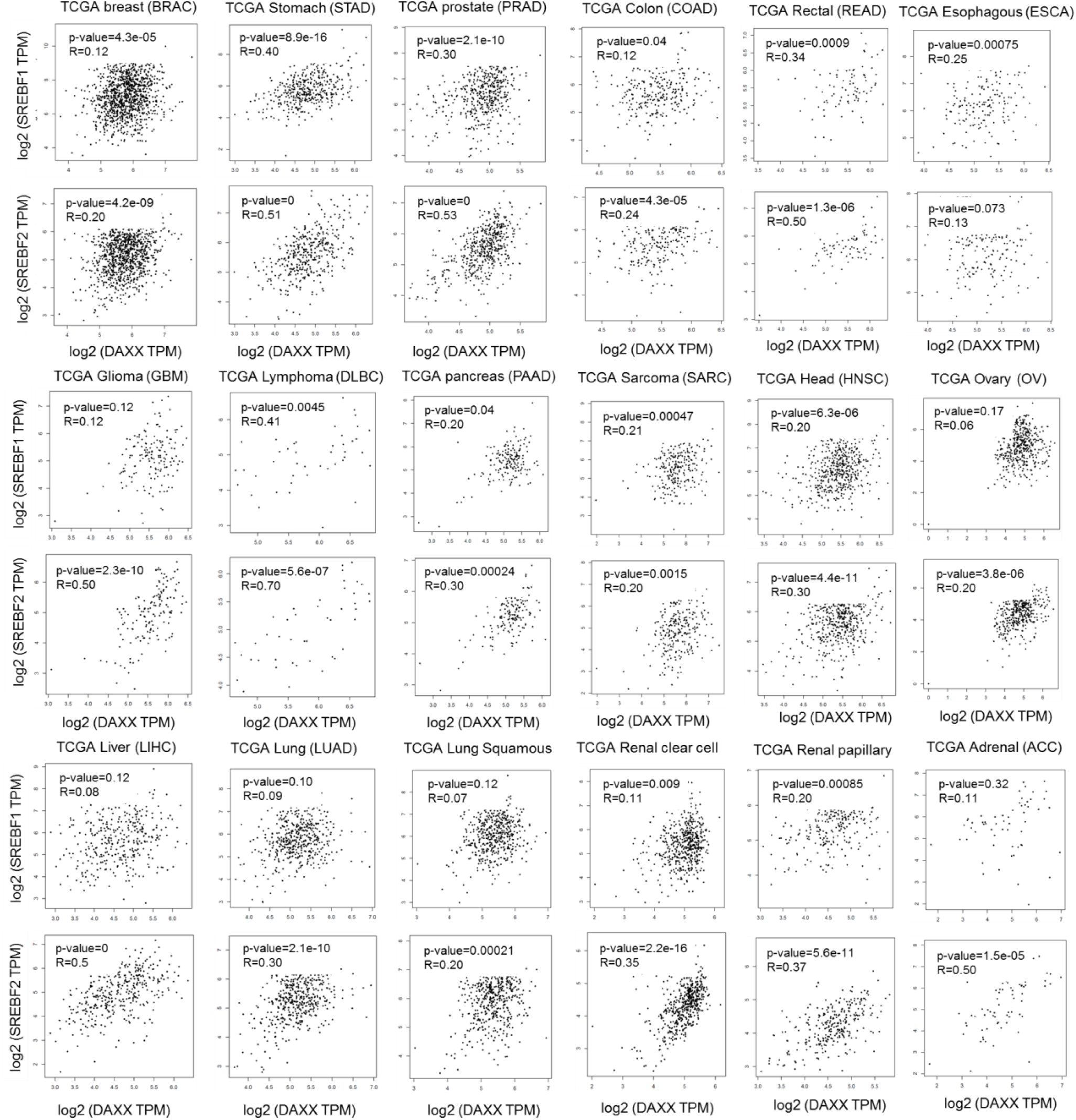
**The mRNA expression of DAXX and SREBP1/2 is positively correlated in eighteen different human cancer types based on the Pearson correlation coefficient analysis of TCGA datasets**.

**Supplementary Figure S5.**
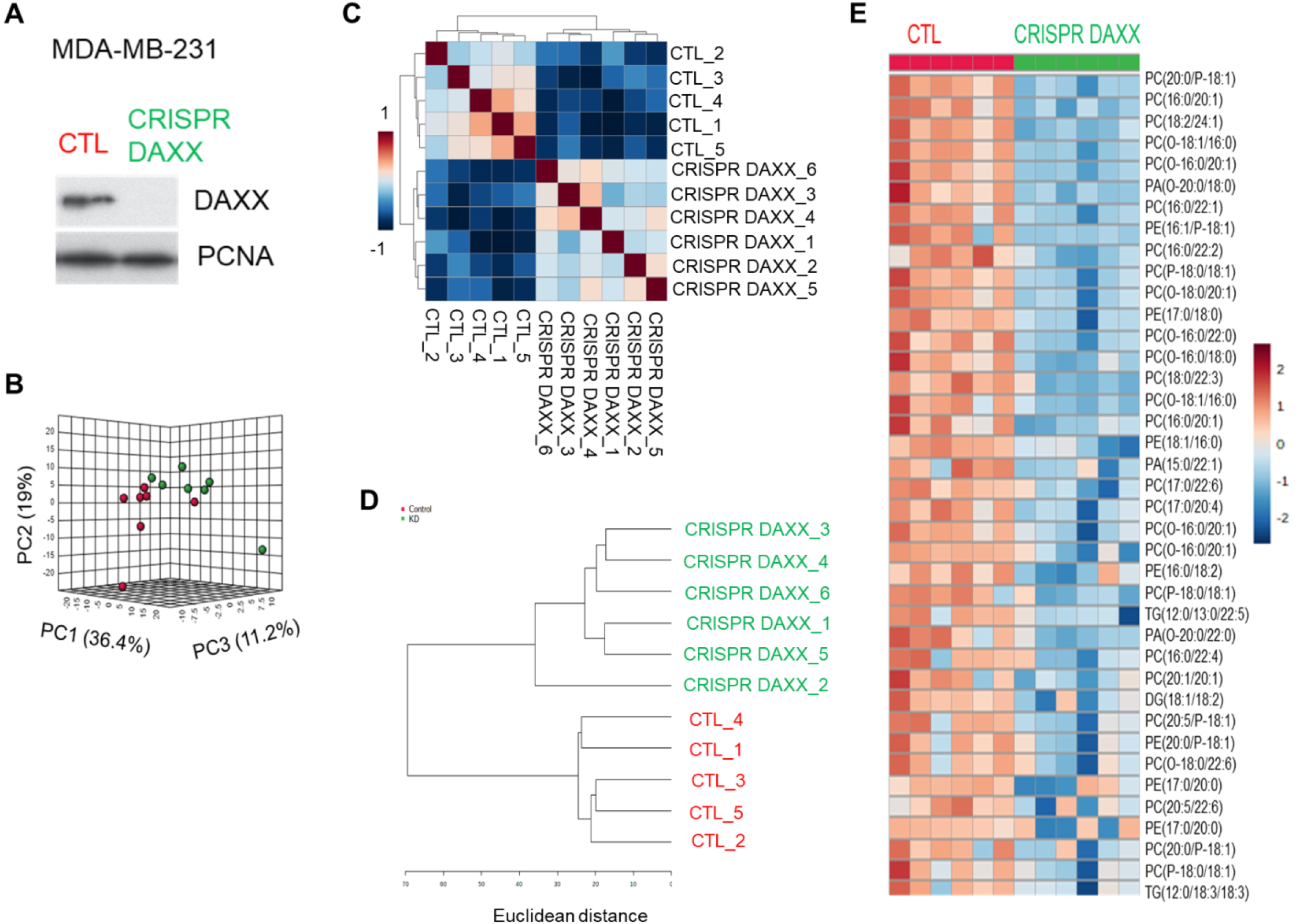
CRISPR/Cas9-mediated DAXX depletion suppressed de novo lipogenesis. (A) Western blot showing DAXX depletion in an MDA-MB-231 clone with a guide RNA targeting DAXX compared to control (CTL) cells. (B) The PCA analysis of lipidomes in control (CTL, red dots) and CRISPR-DAXX MDA-MB-231 (green dots) cells (n=6), which indicates distinct global lipid profiles in these two MDA-MB-231 cell lines. (C) The Pearson correlation coefficient analysis of lipids in CTL and CRISPR-DAXX MDA-MB-231 cells show clear clustering of CTL and CRISPR-DAXX cells. (D) The hierarchical clustering dendrogram analysis of lipids in CTL and CRISPR-DAXX MDA-MB-231 cells. (E) The hierarchical heatmap analysis of lipids in CTL and CRISPR-DAXX MDA-MB-231 cells demonstrate reduced levels of specific lipid molecules in cells with DAXX depletion.

**Supplementary Figure S6.**
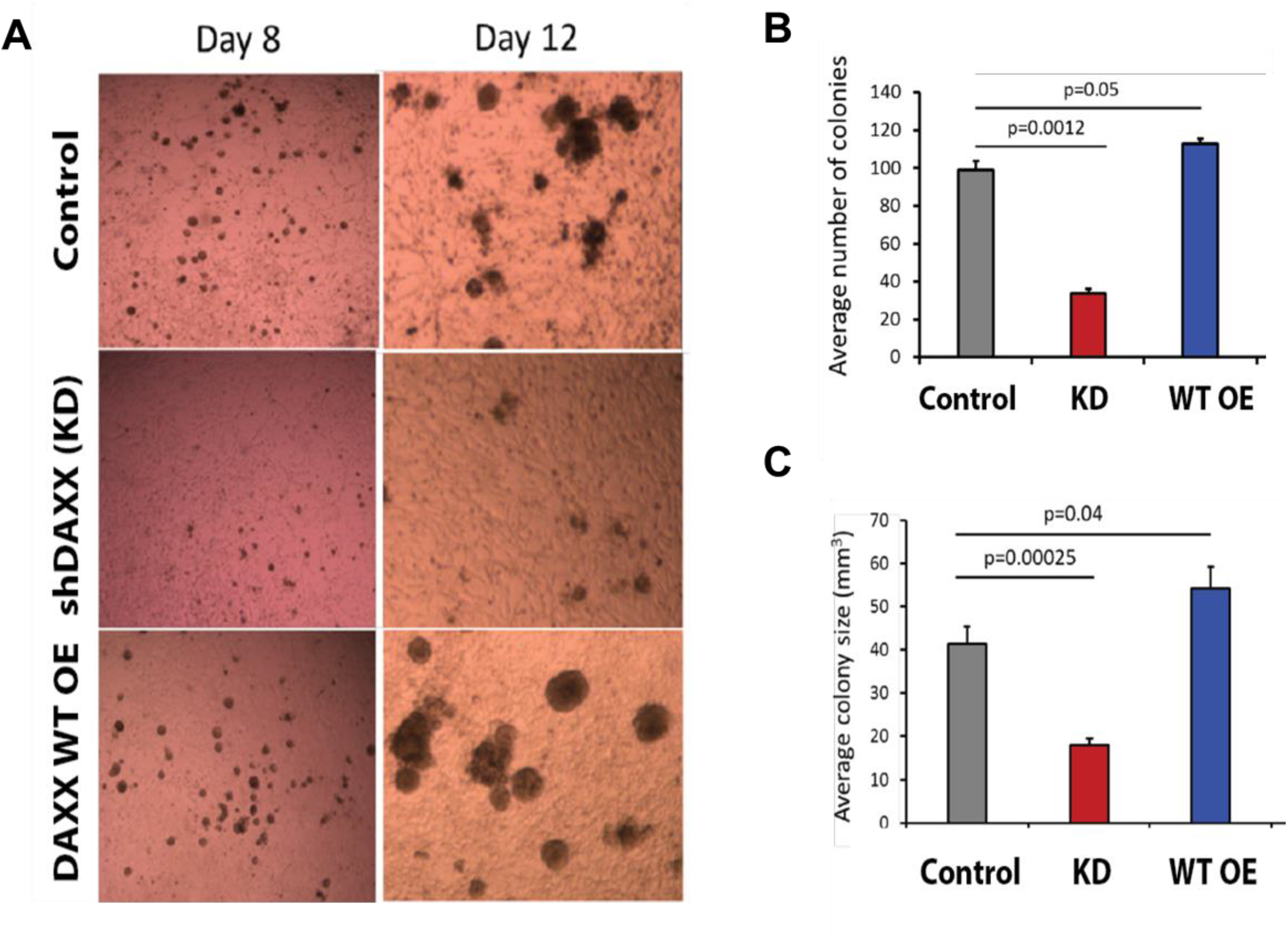
DAXX promotes cell proliferation and 3D colony growth in vitro. (A) MDA-MB-231-derived cell lines (Control, DAXX KD, and DAXX OE) were cultured in a suspension with Matrigel and complete DMEM medium. The 3D colonies of each line were imaged at the indicated time. (B) Average colony number shown as bar graph and were quantified at Day 12. (C) Average colony size shown as bar graph and were quantified at Day 12.

**Supplementary Figure S7.**
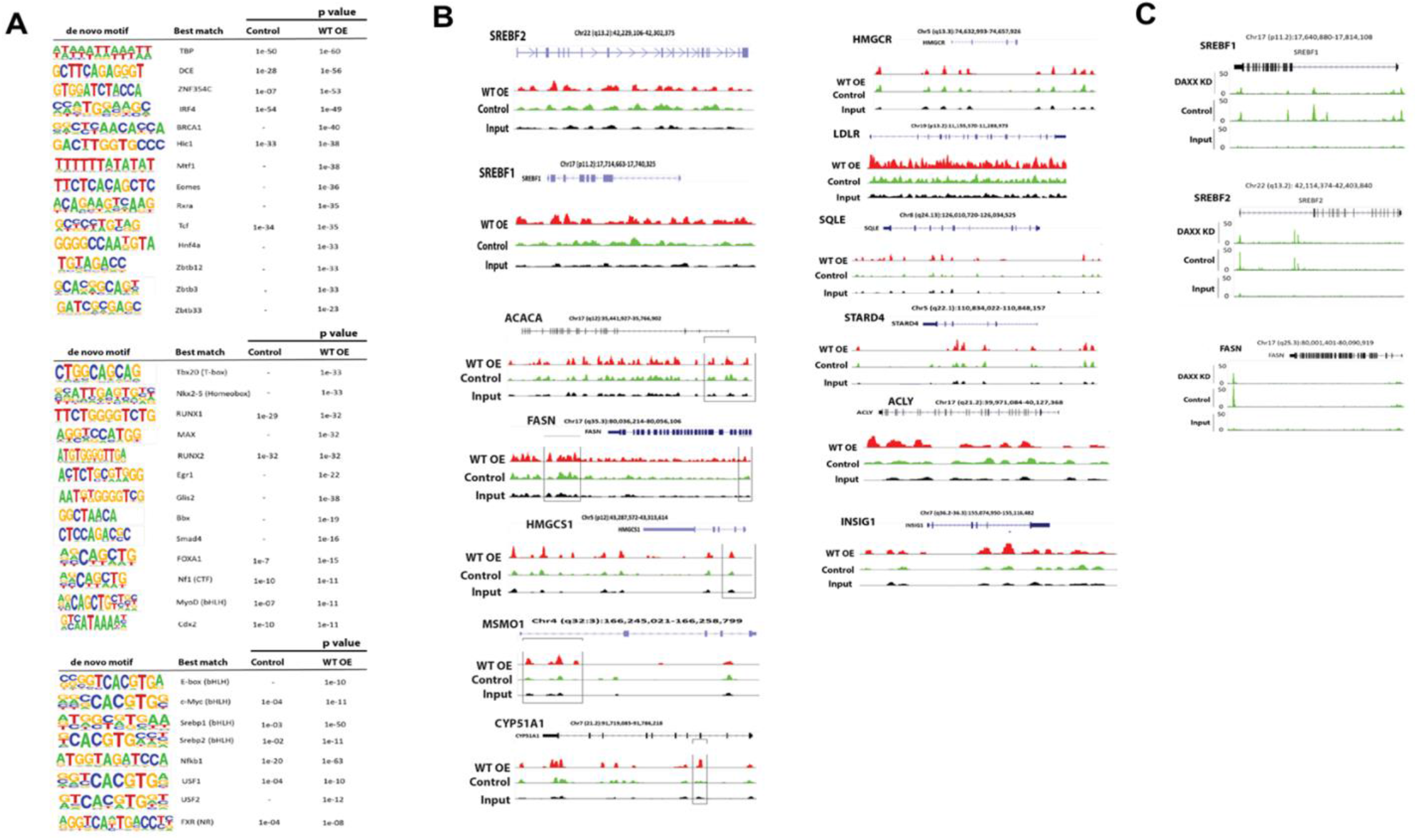
Chromatin-binding activity of DAXX. (A) De novo motifs associated with DAXX as revealed by ChIP-seq. DAXX ChIP-seq and motif analysis were done as in Figure 5. Motifs enriched in MDA-MB-231-derived cells (control and DAXX OE) are shown. (B) DAXX chromatin-binding profiles of the indicated individual lipogenic genes in MDA-MB-231-derived cells (control and wt OE) are depicted. (C) DAXX chromatin-binding profiles of the indicated individual lipogenic genes in PC3-derived cells (control vs. KD; Puto et al., 2015) are shown.

**Supplementary Figure S8.**
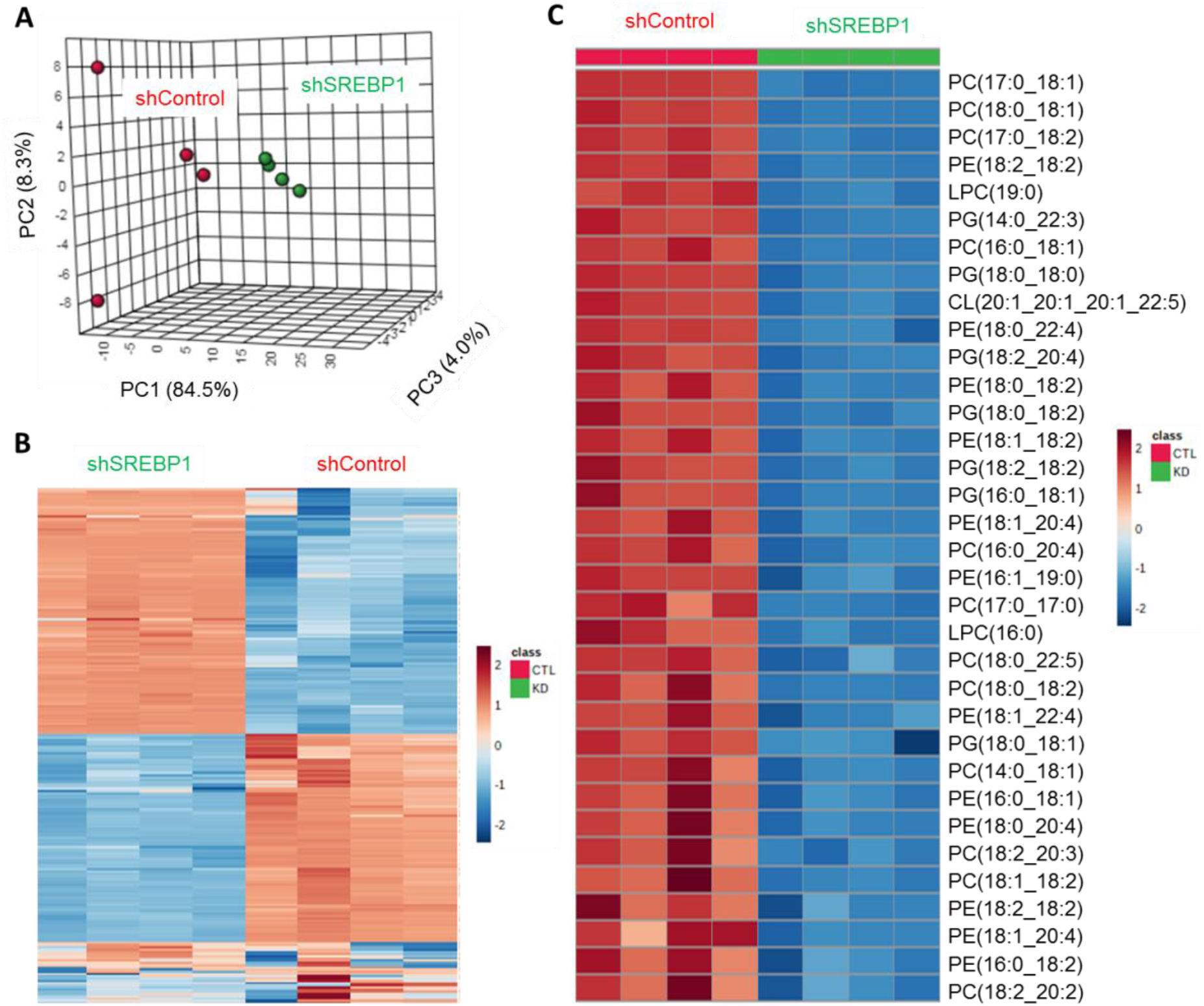
Genetic knockdown of SREBP1 markedly impact global lipid profile. (A) Principal component analysis comparing lipidomes between shControl and shSREBP1 cells derived from MDA-MB-231 cell line. Each dot represents an independent sample (n=4). (B) Hierarchical clustering heatmap analysis demonstrates global lipid landscape in shSREBP1 cells compared to shControl cells. (C) Hierarchical clustering heatmap analysis of top differentially changed lipids including glycerolipid and glycerophospholipid molecules in MDA-MB-231 with SREBP1 KD compared to control cells.

**Supplementary Table S1.**
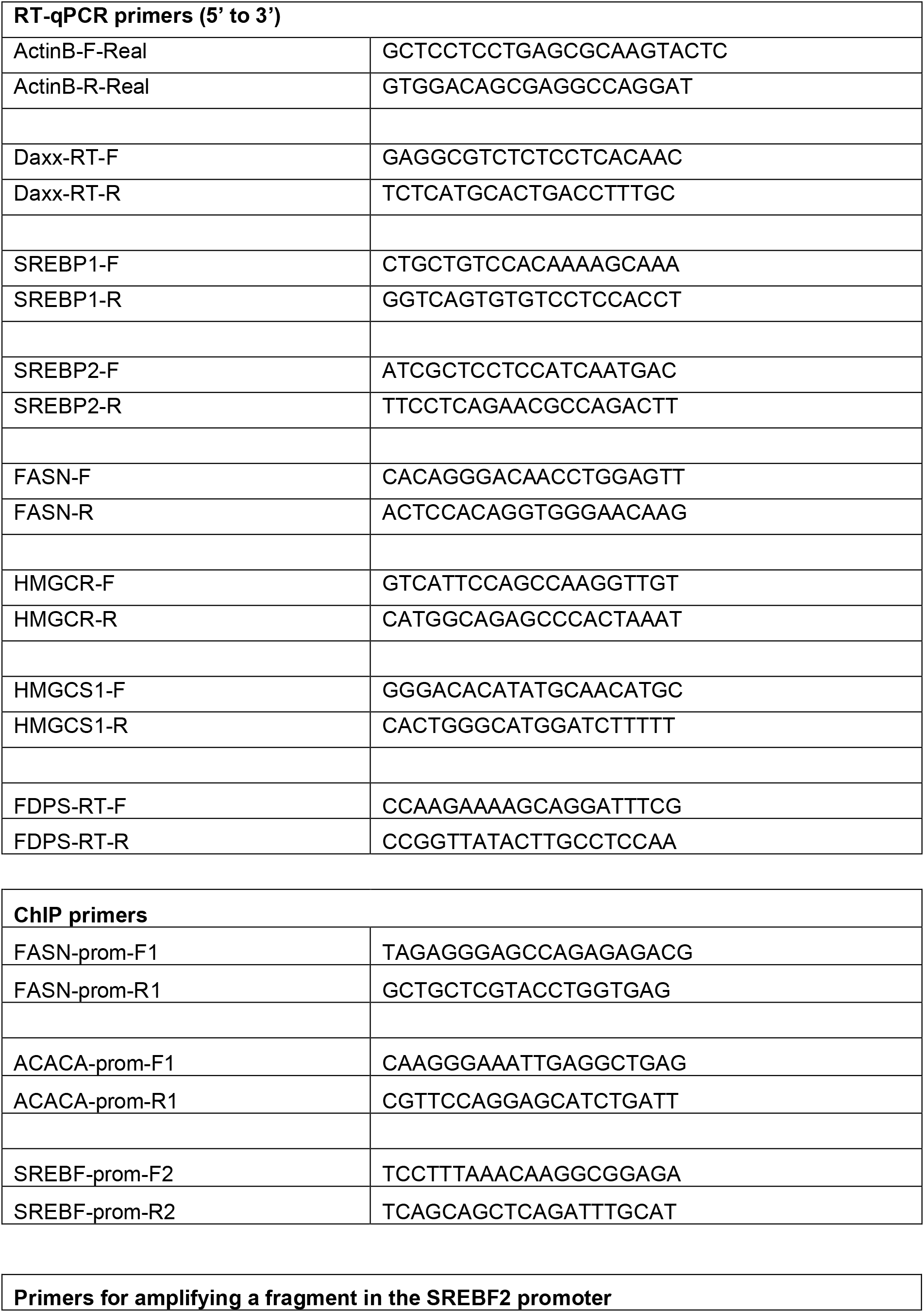
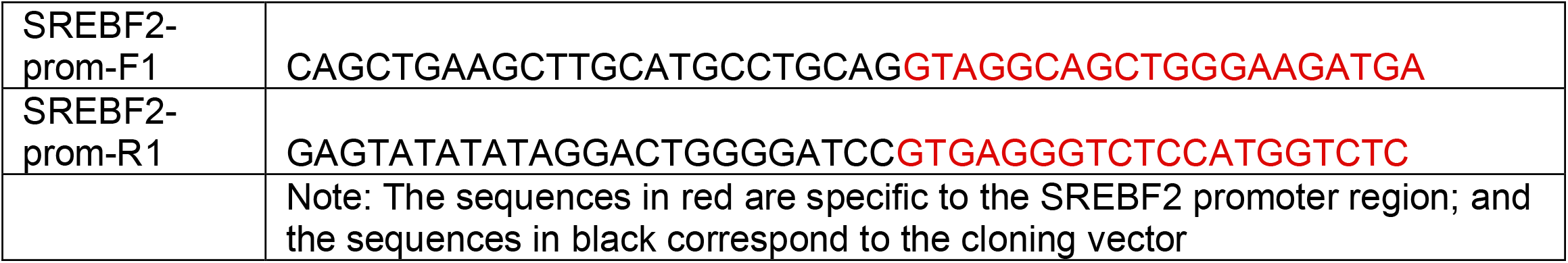
PCR primers used for this study.

**Supplementary Table S2.** Antibodies used in this study.

| Antibody target | Vendor/source | Vendor catalog # | Dilution |
| --- | --- | --- | --- |
| DAXX | Bethyl laboratories | A301-353A | 1:20,000 (IB) |
| DAXX (for IP, IB, IF PLA, and ChIP) | The Developmental Studies Hybridoma Bank | PCRP-DAXX-5G11 | Hybridoma supernatant. 1:100 for IB, 1:5 for IF, and 1:2 for PLA |
| DAXX (for IP and IB) | GenScript, rabbit polyclonal | This study | 1:10,000 for IB |
| FASN (for IB and IF) | ProteinTech | 10624-2-AP | 1:20,000 for IB, and 1:700 for IF |
| FASN | Santa Cruz | SC-55580 | 1:20,000 (IB) |
| ACC1 | Cell Signaling Technology | 3676 | 1:20,000 (IB) |
| ACSS2 | Cell Signaling Technology | 3658 | 1:20,000 (IB) |
| SREBP2 | Abcam | ab30682 | 1:10,000 for IB, 1:300 for IF and 1:100 for PLA |
| SREBP2 | BD Biosciences | 557037 | 1:1,000 (IB) |
| SREBP1 | Santa Cruz | SC-13551 | 1:5,000 (IB) |
| SREBP1 | ProteinTech | 4088-1-AP | 1:3,000 for IB, 1:300 for IF and 1:100 for PLA |
| FLAG | Cell Signaling Technology | 14793 | 1:10,000 (IB) |
| FLAG (IB, IP and ChIP) | Millipore-Sigma | F1804 | 1:1,000 (IB) |
| GFP | Cell Signaling Technology | 2956 | 1:3,000 (IB) |
| PCNA | Epitomics | 2714-1 | 1:20,000 (IB) |
| alpha-Tubulin | Millipore-Sigma | T5168 | 1:50,000 (IB) |
| HSP60 | BD Transduction Laboratories | H99020 | 1:50,000 (IB) |
| Rabbit IgG HRP-linked antibody | Cell Signaling Technology | 7074 | 1:10,000 (IB) |
| Mouse IgG HRP-linked antibody | Cell Signaling Technology | 7076 | 1:10,000 (IB) |
| Normal mouse IgG (for IP/ChIP control) | Santa Cruz | SC-2025 |  |

